# Improvement of affinity and potency of a monoclonal antibody against *Shigella flexneri* 3a O-antigen via phage display and whole-cell in-solution panning

**DOI:** 10.1101/2025.09.11.675504

**Authors:** Nicholas L. Xerri, Sophia Pulido, Mateusz Kędzior, Paul Savarino, Torrey Williams, Robert M. Gallant, Robert W. Kaminski, Devin Sok, Hayden R. Schmidt

**Affiliations:** IAVI Neutralizing Antibody Center, San Diego, California, USA; Latham Biopharm Group, Cambridge, Massachusetts, USA; Global Health Investment Corporation, New York, New York, USA

## Abstract

As rates of antimicrobial resistance (AMR) among bacterial pathogens continue to rise, the discovery and development of novel classes of therapeutics that can serve as alternatives or adjuncts to traditional small-molecule antibiotics, such as monoclonal antibodies (mAbs), is a public health priority. Some of the most promising antigen targets for antibacterial mAbs are surface polysaccharides such as O-antigen (O-Ag), a component of the lipopolysaccharide found on the outer membrane of gram-negative bacteria. However, developing mAbs against bacterial surface polysaccharides with sufficient breadth and potency to be clinically viable is difficult in part because antibodies against polysaccharides are generally low affinity, and the challenging biochemistry of polysaccharides often precludes further affinity maturation of mAbs against these targets *in vitro*. Here, we use a phage display library and a whole-cell in-solution panning strategy to successfully improve the affinity of a mAb against *Shigella flexneri* 3a O-Ag *in vitro* without requiring the purification of the target antigen. We demonstrate that a single mutation can improve apparent affinity approximately 10-fold without detectably increasing polyreactivity, and increased affinity correlates with enhanced potency in antibacterial effector function and anti-virulence assays. In addition, the most potent variants also gained increased breadth, successfully coordinating complement deposition and complement-independent opsonophagocytosis against *S. flexneri* 3b, a serotype weakly recognized by the parent mAb. Altogether, this work represents an important first step towards expanding the antibody engineering toolkit for bacterial surface polysaccharides, which will aid the development of novel mAb therapeutics against AMR bacterial pathogens.

## Introduction

Antimicrobial resistance (AMR) among pathogenic bacteria is a rapidly growing threat that necessitates the development of novel therapeutics to supplement existing antibiotics(1). Monoclonal antibodies (mAbs) may be complementary to antibiotics because they can prevent and treat AMR infections by both recruiting effector cells to kill bacteria via Fc-mediated effector functions (i.e., complement-driven bactericidal activity or complement independent opsonophagocytosis), and by directly interfering with bacterial virulence(2). Some of the most promising antigens for antibacterial immunotherapies are polysaccharides on the bacterial surface. Surface polysaccharides are well validated targets for approved bacterial vaccines (e.g., those against *Haemophilus influenzae* b, *Streptococcus pneumoniae*, *Neisseria meningitidis*, and *Salmonella typhi*) (3–7). This provides a strong rationale for developing anti-polysaccharide mAbs, and anti-polysaccharide mAbs against several bacterial species have been shown to be effective in preclinical animal models (8–17). However, to date, few of these mAbs against bacterial polysaccharides have been evaluated in clinical trials. Panobacumab, an IgM mAb against O-antigen (O-Ag) from serotype O11 *Pseudomonas aeruginosa* obtained promising results in a phase IIa trial among patients with pneumonia, though the effect size was modest and no control arm was present (18), so additional studies are required to fully assess clinical benefit. Gremubamab, a bispecific mAb targeting the exopolysaccharide PsI and the protein PcrV, has entered the clinic (19) and recently shown promise in treating *P. aeruginosa* infections in patients with bronchiectasis, though similarly, this was a proof-of-concept trial and more studies are required to ensure reproducible clinical efficacy (20).

One reason for the small number of both clinical trials and anti-polysaccharide mAb candidates in the pipeline is that discovering and engineering anti-polysaccharide mAbs that have sufficient breadth and potency to be clinically viable is challenging. Breadth for a given anti-polysaccharide mAb is often too narrow for clinical use because the chemical structures of most surface polysaccharides are highly variable (even within the same bacterial species (21). Bacterial strains of a given species are typically grouped into serotypes (often dozens or more) based on the specific chemical structure of key surface polysaccharides (such as the O-Ag, which is a component of gram-negative lipopolysaccharide), and antibody responses against these bacteria are usually serotype-specific (22,23). In addition to breadth, potency is another challenge for anti-polysaccharide mAbs since B cells targeting repetitive polysaccharide antigens undergo T cell independent responses that generate short-lived plasma antibody responses, which are predominately IgM with minimal somatic mutation and low affinity for the target antigen (24). While there are examples of high affinity anti-bacterial polysaccharide mAbs that have undergone extensive affinity maturation isolated from the peripheral blood mononuclear cells of convalescent donors (16), it is unclear how widespread this phenomenon is. In principle, additional high-potency mAbs against bacterial surface polysaccharides can be developed by engineering high-affinity variants of known anti-bacterial polysaccharide mAbs through *in vitro* affinity maturation. The chemical heterogeneity, inherent complexity of polysaccharide biochemistry, and the conformational flexibility of these polysaccharides (leading to a high entropic cost for binding for mAbs (25)) makes achieving *in vitro* affinity maturation of these mAbs is difficult using standard approaches.

To date, relatively few successful attempts at *in vitro* affinity maturation of anti-glycan mAbs have been reported (26–35). Most of these efforts have targeted glycans that are easier to purify and lack the variable, highly repetitive architecture of polysaccharides like O-Ag or capsular polysaccharide. To our knowledge, the only bacterial O-Ag or capsular polysaccharide for which successful *in vitro* affinity maturation has been reported is the O-Ag from *Salmonella* serogroup B. However, the impact that improving the affinity of these mAbs had on antibacterial effector functions was not explored (32,33,36). As a result, the relationship between antibody binding affinity and the potency of specific effector functions is less well characterized for mAbs against bacterial polysaccharides than for those against mammalian glycans.

Here, we utilize a creative whole-cell in-solution phage display panning strategy, coupled with next-generation sequencing, to overcome these challenges and improve the affinity of a prototypical anti-O-Ag mAb specific for *Shigella flexneri* serotype 3a. We demonstrate that one to two mutations are sufficient to not only improve potency across effector functions against the target serotype, *S. flexneri* 3a, but to also expand breadth to serotype 3b. Importantly, a lack of polyreactivity against other bacterial strains or common polyreactivity reagents such as single stranded DNA (ssDNA) indicates that these improvements in binding are specific. Interestingly, binding affinity and potency are generally correlated for most effector functions, but are not necessarily interdependent, particularly for complement-independent opsonophagocytosis. Altogether, these data provide a framework for affinity maturing mAbs against bacterial polysaccharides *in vitro* and demonstrate that (at least in this test case) few mutations are needed to significantly improve the breadth and potency of an anti-O-Ag antibody.

## Results

### hFlex3a2 functions as a chimeric huIgG1, but not as an scFv

hFlex3a2 is a murine hybridoma IgG3 that binds to *S. flexneri* 3a O-Ag and can coordinate complement-mediated bacteriolysis of this serotype in an antibody bactericidal assay (37). We determined the sequence of this mAb (Table S1) and found that it has many of the properties commonly associated with anti-O-Ag mAbs. The putative variable gene segment of the heavy chain of hFlex3a2 differs from the germline segment IGHV1-4*02, its closest germline progenitor, by only four amino acids, while the putative variable gene segment of the light chain of hFlex3a2 is entirely identical to its closest germline segment IGKV5-39*01 (Figure 1A). The high degree of similarity to its closest predicted germline progenitor suggests that immunizing mice with purified lipopolysaccharide (LPS) did not stimulate high levels of somatic mutation *in vivo.* The putative VH and VL genes were grafted onto the human IgG1 constant regions to create a chimeric huIgG1 and this antibody was confirmed by ELISA to bind to *S. flexneri* 3a outer membrane vesicles (OMVs) (Figure 1B), a proxy for the bacterial cell surface (38), as well as to coordinate complement-mediated bacteriolysis of *S. flexneri* 3a in a luminescent antibody bactericidal assay (L-ABA, Figure 1C). Additionally, hFlex3a2 was highly serotype-specific, binding to only the surface of *S. flexneri* 3a that expresses wildtype O-Ag, when screened against a panel of serotypes and an O-Ag knockout mutant (Figures S6A and 1D). Thus, the low level of somatic mutation, combined with its narrow breadth, make hFlex3a2 representative of a canonical mAb against a bacterial surface polysaccharide.

**Fig 1:**
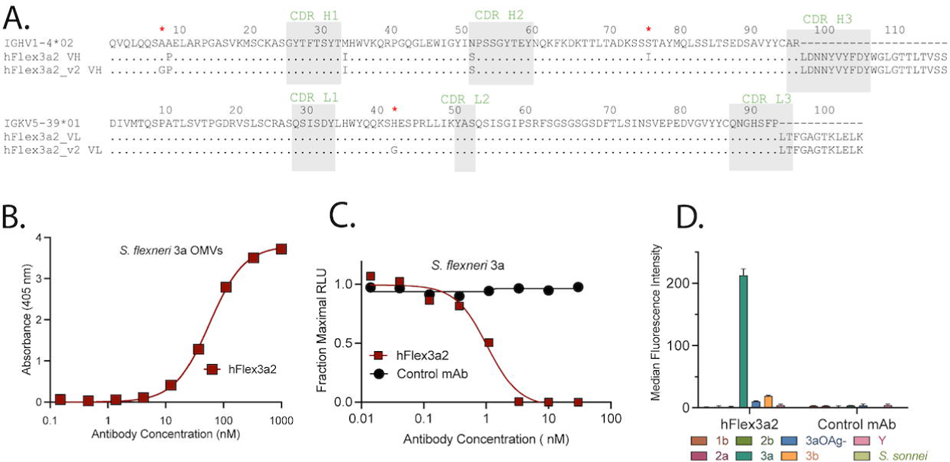
Analysis of binding and function of putative hFlex3a2 as a huIgG1 chimera. (A) Alignment of putative hFlex3a2 VH and VL protein sequences to their closest germline progenitors shows high levels of similarity. Residues that were modified to create mAb hFlex3a2_v2 are denoted by (*). Alignment and CDR predictions were determined with IMGT/V-Quest and rare residues were identified with abYsis (45,46). (B) Titration ELISA measures the binding of increasing concentrations of hFlex3a2 huIgG1 to 2.5 μg/mL of *S. flexneri* 3a OMVs. This experiment is the average of three independent replicates each performed in duplicate. (C) L-ABA measuring the ability of increasing concentrations of hFlex3a2 huIgG1 and a non-binding control mAb, to coordinate complement-mediated bacterial killing of *S. flexneri* 3a. L-ABA is the average of three biological replicates performed in duplicate. (D) Surface staining with 50 nM of mAb shows that hFlex3a2, and not a non-binding control mAb, binds specifically to *S. flexneri* 3a wildtype O-Ag. This experiment is the average of three experimental replicates. Error bars represent standard deviation.

We sought to improve the affinity of hFlex3a2 against *S. flexneri* 3a O-Ag using phage display. Since hFlex3a2 would be displayed as an scFv, we wanted to confirm that the antibody retains functionality in this format. However, binding of *S. flexneri* 3a OMVs by hFlex3a2 in an scFv format could not be detected by ELISA (Figure 2A). This is not surprising, as scFvs often exhibit lower target affinity and are less stable than their IgG counterparts (39,40). In an attempt to restore function to hFlex3a2 as an scFv, we identified two residues in the framework region of the VH and one in the VL that were rare and thus may be potentially problematic (41). We mutated all three of these residues to the amino acid most frequently found at their respective positions when compared to a database of other murine antibodies. We called this new mAb hFlex3a2_v2 (Figure 1A). Two of these new mutations were away from the closest germline sequence, while the other reverted that position back to its predicted germline amino acid (Figure 1A). As an scFv, binding of hFlex3a2_v2 to *S. flexneri* 3a OMVs, but not to the unrelated antigen ovalbumin, was observed (Figure 2A). Antigen binding was further confirmed by surface staining, which demonstrated that hFlex3a2_v2 scFv, but not the original hFlex3a2 scFv binds to the surface of *S. flexneri* 3a (Figure 2B). Importantly, as a chimeric huIgG1, hFlex3a2_v2 exhibited antigen binding similar to the parent mAb, demonstrating that the rare residues we identified did not impact IgG antibody binding (Figure 2C). Since hFlex3a2_v2 retained binding as a huIgG1, and restored binding as an scFv, while retaining antigen specificity, affinity maturation by phage display was completed using the hFlex3a2_v2 background.

**Fig 2:**
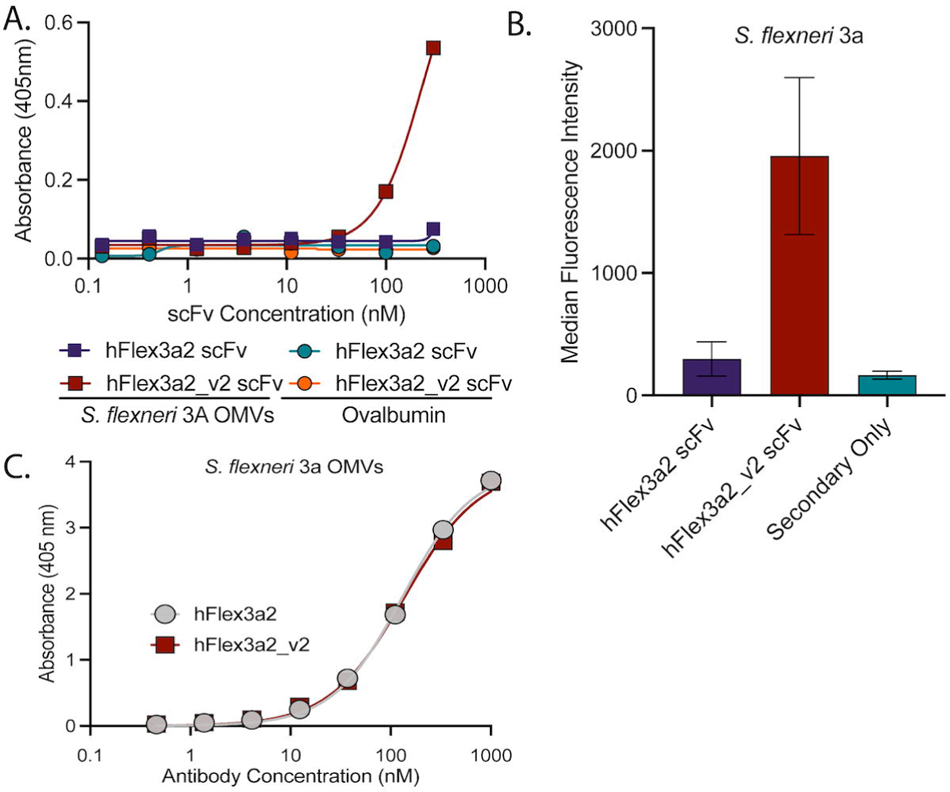
Analysis of binding of hFlex3a2 and hFlex3a2_v2 as an scFv. (A) Titration ELISA shows that at concentrations above 100 nM of scFv, hFlex3a2_v2, but not hFlex3a2, binds to *S. flexneri* 3a OMVs (2.5 μg/mL). Neither scFv bound to the unrelated antigen ovalbumin (5 μg/mL). This experiment was repeated three times, and a representative experiment is shown. (B) Titration ELISA of hFlex3a2_v2 reformatted as a chimeric huIgG1 and the original hFlex3a2 huIgG1 shows similar binding to *S. flexneri* 3a OMVs (2.5 μg/mL). This experiment is the result of two independent replicates. (C) Surface staining confirms that in the scFv format, hFlex3a2_v2, but not hFlex3a2, binds *S. flexneri* 3a. scFvs were tested at a concentration of 1μM. This experiment represents two replicates. Error bars represent standard deviation.

### Whole-cell in-solution panning enriched for mutations in the VL

To test whether phage display can be used to identify single mutations that improve hFlex3a2_v2 affinity, we designed two double-barcoded site-saturated mutagenesis scanning libraries: one for VH and one for VL. Each library contained single amino acid substitutions besides a cysteine at every site of either the VH or VL, leading to total library sizes of 2,223 and 1,995 variants for the VH and VL libraries respectively. Flanking synonymous codons were used to barcode mutation sites, enabling bioinformatic identification of sampled positions for mutation enrichment analysis. Initial efforts to fix purified *S. flexneri* 3a lipopolysaccharide to a plate for selection via solid-phase panning were unsuccessful. Whole-cell in-solution panning was previously used to discover antibodies that bind to bacterial surface carbohydrates (42), and so we adopted a similar panning strategy for affinity maturing our existing O-Ag antibody (Figure 3A). The phage display libraries were panned in solution for two rounds against either the target bacterium *S. flexneri* 3a or *S. sonnei*, which is a closely related bacterium with an entirely different O-Ag structure (43). Input and output page libraries were subsequently sequenced by next-generation sequencing. We reasoned that variants enriched specifically after panning against *S. flexneri 3a*, but not *S. sonnei*, were likely to exhibit improved binding to the target relative to the parental mAb, and these variants could be evaluated further. This approach also had some advantages over other panning strategies. Specifically, the approach did not require large-scale purification of O-Ag, and the approach also allowed for panning on O-Ag in its native conformation and abundance without any biochemical modifications.

**Fig 3:**
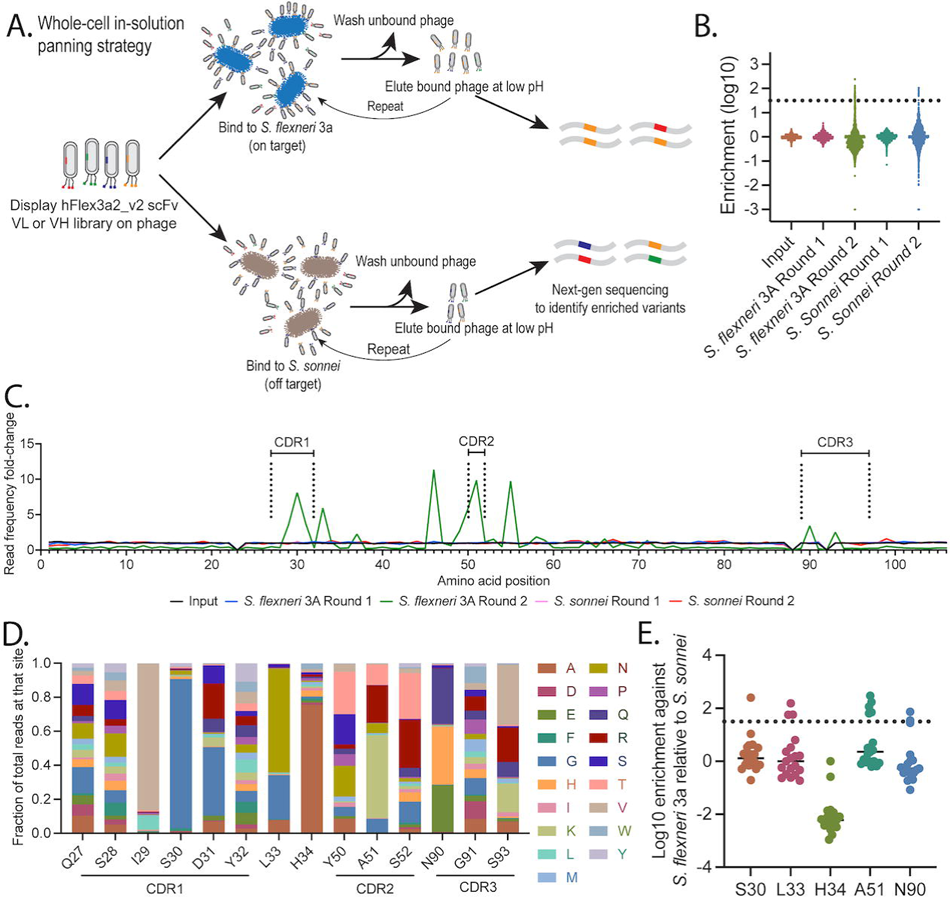
Whole-cell in-solution panning enriches for VL variants. (A) Whole-cell in-solution panning strategy. To identify better binding variants of hFlex3a2_v2 scFv, we panned a phage-displayed site-saturated mutagenesis scanning library of VL or VH against either *S. flexneri* 3a or *S. sonnei* in-solution for two rounds and sequenced output phage by next-generation sequencing to identify variants that were specifically enriched against *S. flexneri* 3a. (B) log10 enrichment of variants in the VL library after two rounds of panning against *S. flexneri* 3a and *S. sonnei*. Dotted line is 20 standard deviations above the mean mutation enrichment in the input library and indicates a stringent threshold set to distinguish bacteria-mediated enrichment from background variability. (C) Fold-change in the frequency of next-generation sequencing reads for variant mutations at each site on hFlex3a2_v2 VL after panning, relative to their frequency in the initial library, shows that certain residues that cluster in and around the CDRs have a large increase in read frequency. (D) The percent distribution of all variants for each site of the CDRs after two rounds of panning against *S. flexneri* 3a shows that at some sites (I29, S30, H34) a single variant, or at other sites (L33, A51, N90, S93) a handful of variants were enriched for. (E) Specificity of mutational enrichment was determined. Each dot represents the enrichment of a variant against *S. flexneri* 3a relative to *S. sonnei* at the indicated residue after two rounds of panning. Positive numbers indicate variants that were enriched against *S. flexneri* 3a relative to *S. sonnei*, zero indicates similar enrichment against both bacterial species, and negative numbers indicate variants that were enriched for against *S. sonnei* relative to *S. flexneri* 3a. Dotted line is 20 standard deviations above the mean mutation enrichment in the input library.

After two rounds of panning with the VH library, no enrichment of hFlex3a2_v2 variants was detected (Figure S1A and S1B). In contrast, specific enrichment did occur for variants in the VL library. Despite all the variants being of roughly equal abundance in the input library, by the second round of panning, positively and negatively enriched variants were identified when panned against *S. flexneri* 3a as well as *S. sonnei* (Figure 3B). After two rounds of panning against *S. flexneri* 3a, a strong increase in the frequency of next-generation sequencing reads for variant mutations mapping to residues that cluster around the VL CDRs was observed (Figure 3C). This enrichment was driven by the accumulation of reads mapping to either a single variant or a subset of variants at these sites (Figure S2). In some cases, such as at residue S30, a single variant was found to dominate relative to other variants with mutations at that site, while at other residues, such as N90, a few variants were found to be similarly abundant (Figure 3D). To determine whether highly enriched variants identified during phage display screening were of higher affinity than the parental antibody, we picked four variants — S30, L33, A51, and N90 — for further investigation. We chose these variants because all four had large increases in read frequencies after panning, suggesting that these residues may be important. Additionally, each variant was specifically enriched following panning against *S. flexneri* 3a relative to *S. sonnei* (Figures 3E and S3 and Table 1). Lastly, these variants are located in or adjacent to CDRL1 (S30 and L33), CDRL2 (A51) or CDRL3 (N90). Since no variants in the VH were enriched for, we will refer to the hFlex3a2_v2 variants solely by their VL mutation(s) from here on.

### Variants enriched in phage display improve mAb binding to *S. flexneri* 3a O-Ag as chimeric huIgG1s

We hypothesized that the selected variants of hFlex3a2_v2 that were highly enriched against *S. flexneri* 3a but not *S. sonnei* (S30G, L33N, A51R, and N90H) were likely to have genuinely higher affinity against *S. flexneri* 3a O-Ag relative to the parent mAb. For a comparison, we chose one additional variant — H34A — that was highly enriched against both *S. flexneri* 3a and *S. sonnei* (Figures 3D, 3E, S3, and Table 1). All the selected variants were produced as chimeric huIgG1s for further characterization (Table 1).

Surface staining showed that all the variants retained their ability to bind to the surface of *S. flexneri* 3a (Figure 4A). Titration ELISA also confirmed binding to *S. flexneri* 3a OMVs (Figure 4B) and showed that all four of the variants that were specifically enriched against *S. flexneri* 3a (S30G, L33N, A51R, and N90H) had higher apparent affinities than their parental mAb hFlex3a2_v2, while the variant that was enriched against both *S. flexneri* 3a and *S. sonnei* (H34A) had a lower apparent affinity (Figure S4A). The ELISA binding curves were used to calculate EC_50s_ (Table 1 and Figure 4C). The parent mAb hFlex3a2_v2 had an EC_50_ of 57.81 nM. L33N, the variant with the highest apparent affinity, had an EC_50_ of 4.99 nM, while variant H34A had the lowest apparent affinity (EC_50_ 205.09 nM). This is consistent with the hypothesis underlying our panning strategy, that specific enrichment against *S. flexneri* 3a provides a means of identifying variants that improve target affinity.

**Fig 4:**
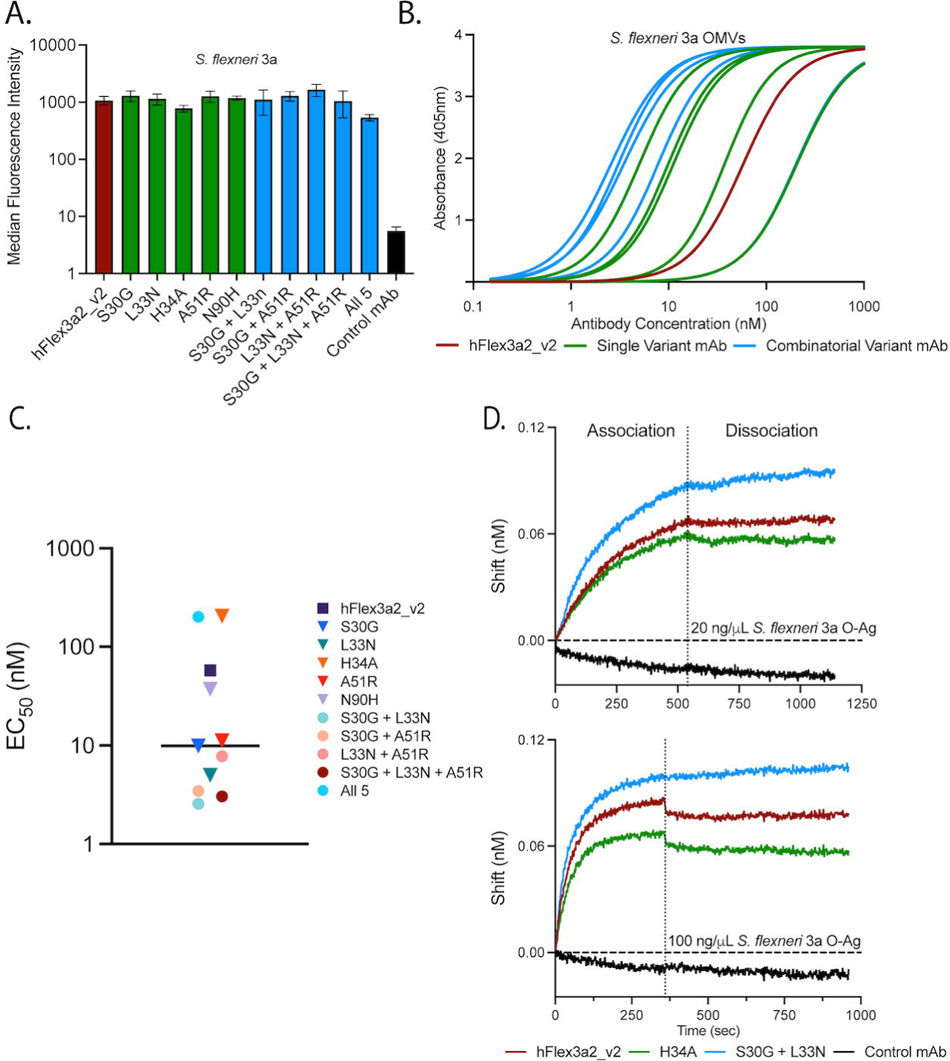
Binding analysis of hFlex3a2_v2 variants specifically enriched against *S. flexneri* 3a and produced as huIgG1 chimeras. (A) Surface staining experiment confirms that 50 nM of the parental mAb (red), single amino acid variants (green), and combinatorial variants (blue) bind *S. flexneri* 3a ∼100-fold higher than a non-binding control mAb (black) as indicated by a median fluorescence intensity. Surface staining is the average of three biological replicates. Error bars represent standard deviation. (B) Binding curves from a titration ELISA against *S. flexneri* 3a OMVs (2.5 μg/mL) show relative binding potencies of the parent mAb (red), single amino acid variants (green), and combinatorial variants (blue). ELISA binding curves are the average of three independent replicates each performed in duplicate (C) EC_50_ values calculated from the ELISA in panel B show relative steady-state apparent affinities of hFlex3a2_v2 variants. Black line indicates the median EC_50_. (D) Binding of hFlex3a2_v2, H34A, S30G + L33N, and a non-binding control as huIgG1s to purified *S. flexneri* 3a O-Ag was assessed by BLI. This experiment is the average of two experiments performed with independently purified batches of mAbs.

We next wanted to determine whether combining the most beneficial single amino acid mutations — S30G, L33N, and A51R — would synergize with one another to further improve the affinity of hFlex3a2_v2. We constructed all the possible double and triple combinatorial mutants. Since all the combinatorial variants retained their ability to bind *S. flexneri* 3a (Figure 4A), we performed ELISA to determine their apparent affinities. All of the combinatorial variants except L33N + A51R had synergistic improvements in apparent affinity, meaning that the combinatorial variant had a lower EC_50_ than that of any single mutation contained in the combination (Table 1, and Figures 4B, 4C, and S4A). Variant S30G + L33N was the most potent binder, with an EC_50_ = 2.56 nM. We also tested a variant combining all 5 of the single mutations and found that its apparent affinity (EC_50_ = 201.35 nM) was similar to that of H34A alone, suggesting that the presence of mutation H34A may be dominantly detrimental over the other beneficial mutations. Due to the oligomeric structure of O-Ag, it is difficult to accurately measure its molar concentration in solution, complicating efforts to precisely measure the binding kinetics between the mAbs and purified O-Ag *in vitro*. However, biolayer interferometry (BLI) confirmed that the parent mAb, and the most potent and least potent variants in ELISA, S30G + L33N and H34A respectively, can all bind purified *S. flexneri* 3a O-Ag (Figure 4D). Consistent with the selected mutations affecting binding behavior, at both concentrations of O-Ag tested, S30G + L33N reached a higher plateau in BLI than the parental mAb, while H34A reached a lower plateau. Lastly, a panel of polyspecificity reagents were used in ELISA to demonstrate that the mAb variants had limited polyreactivity against the antigens tested, confirming that the better binding variants retained their antigen specificity (Figure S4B).

### Improving binding of hFlex3a2_v2 enhances effector and anti-virulence functions

To probe the relationship between improved O-Ag affinity and potency across antibacterial functions, we tested our panel of hFlex3a2 variants in a series of effector function and virulence assays. A Luminescent Antibody Bactericidal Assay (L-ABA), which measures the ability of antibodies to coordinate complement-mediated bacteriolysis, showed that the parent antibody was already potent at this effector function (IC_50_ = 1.86 nM, Table 1, Figures 5A and S5A). Nonetheless, the best binding single amino acid variant, L33N, modestly but reproducibly improved upon the parent mAb (IC_50_= 1.32 nM, p=0.04), while the best binding combinatorial variant, S30G + L33N, had a more substantial ∼2.7-fold improvement in potency (IC_50_ = 0.68 nM, p<0.0001). The worst binding variant H34A had the weakest potency in L-ABA (IC_50_ = 4.86 nM, p<0.0001), showing that binding strength and *in vitro* complement-mediated bacteriolysis are correlated (R^2^=0.91, Figure S5B). Similarly, when hFlex3a2_v2 was tested in a complement-independent opsonophagocytosis assay (OPA) at a 30 nM concentration, 17.7% of monocytes were cell trace violet positive (CTV+), indicating that they had phagocytosed *S. flexneri* 3a that was opsonized with this mAb (Figure 5B). The gating strategy for OPA is shown in Figure S4C. Although hFlex3a2_v2 already had the ability to facilitate OPA activity, L33N, the best binding single amino acid variant, was almost 2-fold more potent at the same concentration (33.7% CTV+, p = 0.02). The best binding combinatorial variant S30G + L33N was the most potent at facilitating OPA (38.6% CTV+, p <0.005). Despite having little improvement in binding and L-ABA compared to the parental mAb, variant N90H was the third most potent in OPA (32.2% CTV+, p = 0.04). Similarly, despite exhibiting lower apparent affinity, H34A and the combination of all 5 mutations had similar potency in OPA to the parent antibody at 30 nM (Figure 5B). This demonstrates that, while largely correlated, other factors beyond binding strength also impact O-Ag mAb function in the OPA.

**Fig 5:**
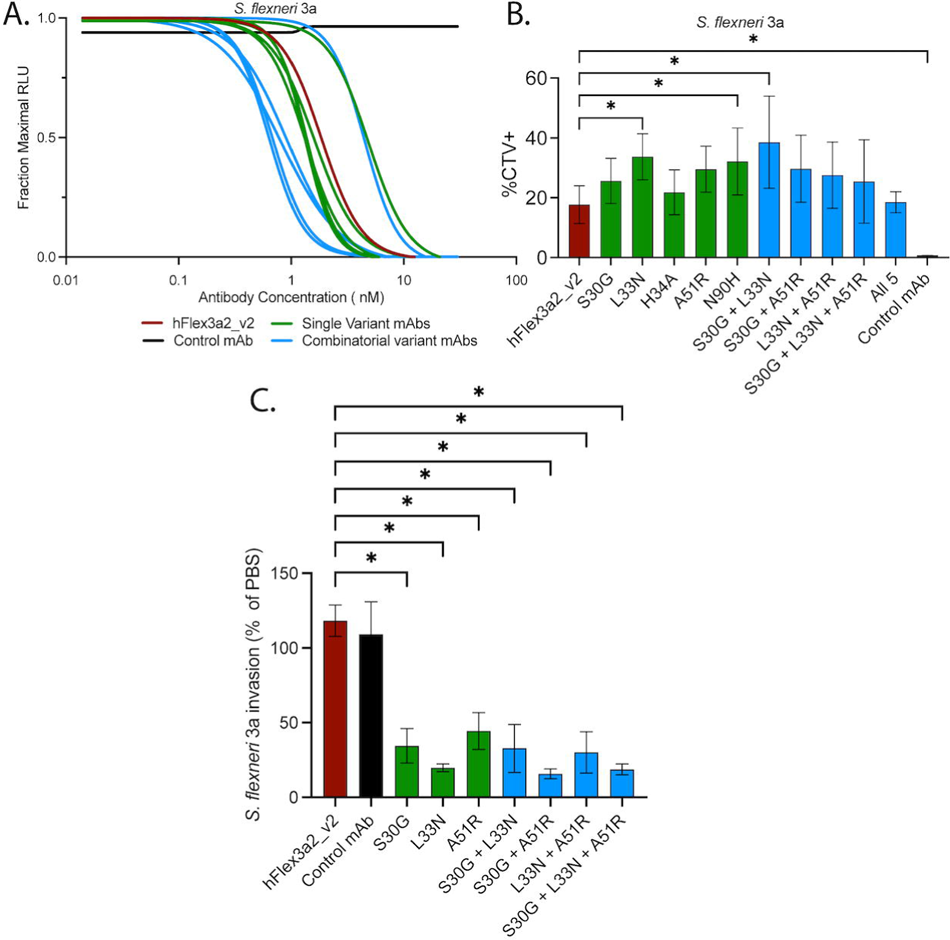
Functional analysis of hFlex3a2_v2 variants against *S. flexneri* 3a. (A) L-ABA assay comparing the ability of hFlex3a2_v2 variants and a non-binding control to coordinate complement-mediated bacteriolysis of *S. flexneri* 3a. L-ABA is the average of three biological replicates each performed in duplicate. p-values were calculated on logIC_50_ values by repeated measure Anova with Dunnett’s correction for multiple comparisons. (B) OPA assay measures the percent of THP-1 cells that have phagocytosed *S. flexneri* 3a that was opsonized with 30 nM of the indicated mAb. This experiment is the result of five biological replicates each performed in duplicate, and p-values were calculated by repeated measure ANOVA with Dunnett’s correction for multiple comparisons. Error bars represent standard deviation. (C) Invasion assay shows the percent invasion of *S. flexneri* 3a incubated with 1μM of the indicated mAb normalized to the invasion rate of *S. flexneri* 3a in PBS. This experiment is the result of four biological replicates each performed in duplicate, and p-values were calculated by repeated measure ANOVA with Dunnett’s correction for multiple comparisons. Errors bars represent standard error of the mean (S.E.M.). For all plots, p values < 0.5 are denoted by a *.

In addition to promoting effector functions that modulate the immune system, mAbs can also directly neutralize bacterial virulence. As an intracellular pathogen, invasion of colonic epithelial cells is an indispensable step in the pathogenesis of *S. flexneri*. An *in vitro* invasion assay showed that, relative to PBS or a non-binding control mAb, hFlex3a2_v2 does not significantly block *S. flexneri* 3a invasion of a monolayer of HeLa cells (Figure 5C). However, the highest affinity single amino acid variants and combinatorial variants all blocked *S. flexneri* 3a invasion by at least 60% (Figure 5C). This demonstrates that for anti-O-Ag antibodies, increasing affinity can help to directly neutralize bacterial virulence.

### Improving affinity of hFlex3a2_v2 against *S. flexneri* 3a increases its functional potency against *S. flexneri* 3b

Lastly, we wanted to determine whether higher affinity variants of hFlex3a2_v2 would also have increased breadth compared to their parent mAb. Although the precise epitope of hFlex3a2_v2 is not known, we reasoned that many of the components of *S. flexneri* 3a O-Ag are also found in the O-Ag of other *S. flexneri* serotypes, and some of the variants may have increased affinity against some O-Ag component common to other *S. flexneri* serotypes. We performed a surface staining experiment with a panel of *S. flexneri* serotypes that all share the same core O-Ag sugars, but with different modifications, as well as *S. sonnei*, which has an entirely unrelated O-Ag composition (Figure 6A). As previously shown, all hFlex3a2_v2 variants bound to *S. flexneri* 3a and binding to a mutant *S. flexneri* 3a O-Ag-strain (Figure S6A) was 10-fold lower than to *S. flexneri* 3a that expresses wildtype O-Ag, confirming that O-Ag is necessary for binding. For all the hFlex3a2 variants tested, no reactivity to *S. flexneri* serotypes 1b, 2a, 2b, Y, or *S. sonnei* was observed (Figure 6B). In contrast, surface staining against *S. flexneri* 3b showed that hFlex3a2_v2 weakly cross-reacts with this serotype. The single amino acid variant L33N and the combinatorial variant S30G + L33N had surface staining median fluorescence intensities that were 4-fold and 5.7-fold higher, respectively.

**Fig 6:**
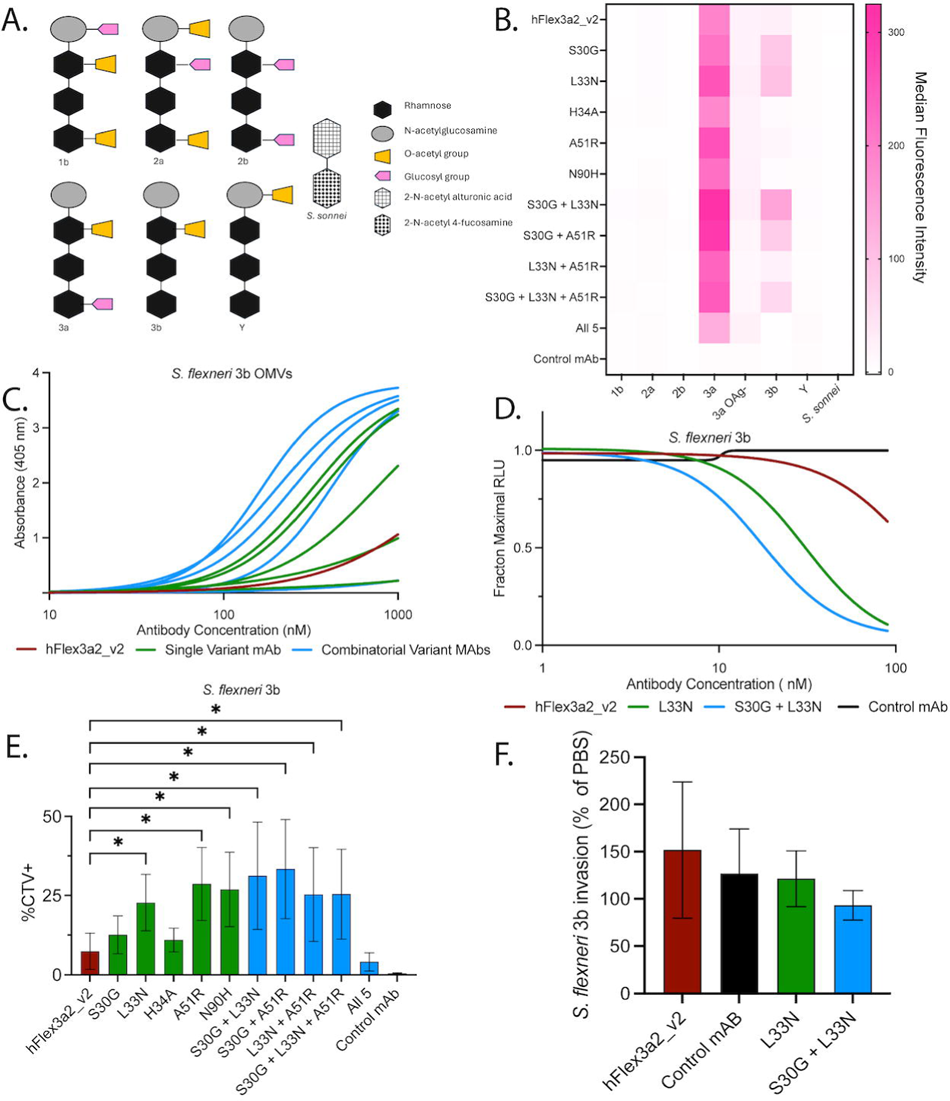
Binding and functional analysis of hFlex3a2_v2 variants against *S. flexneri* **3b.** (A) Schematic depicting the O-Ag structures of our panel of *S. flexneri* serotypes. (B) Surface staining with 50 nM of hFlex3a2_v2 variants and a non-binding control mAb to a panel of *S. flexneri* and *S. sonnei* serotypes. This experiment is the average of two replicates. (C) Binding curves from a titration ELISA against *S. flexneri* 3b OMVs (2.5 μg/mL) show relative binding potency of the parent mAb (red), single amino acid variants (green), and combinatorial variants (blue). ELISA binding curves are the average of two replicates. (D) L-ABA assay comparing the ability of hFlex3a2_v2, L33N, and S30G + L33N to coordinate complement-mediated bacteriolysis of *S. flexneri* 3b. L-ABA is the average of four biological replicates each performed in duplicate. (E) OPA assay measures the percent of THP-1 cells that have phagocytosed *S. flexneri* 3b that was opsonized with 30 nM of the indicated mAb. This experiment is the result of five biological replicates each performed in duplicate, and p-values were calculated by repeated measure ANOVA with Dunnett’s correction for multiple comparisons. Error bars represent standard deviation. (F) Invasion assay shows the percent invasion of *S. flexneri* 3b incubated with 1μM of the indicated mAb normalized to the invasion rate of *S. flexneri* 3b in PBS. This experiment is the result of four biological replicates each performed in duplicate. Error bars represent standard error of the mean (S.E.M.). for all plots, p values < 0.5 are denoted by a *.

Since surface staining experiments showed that the higher affinity variants of hFlex3a2_v2 bind to *S. flexneri* 3b, we wanted to further characterize their effect on binding and effector function. ELISA measuring binding to *S. flexneri* 3b OMVs, in agreement with the surface staining experiment, showed that hFlex3a2 weakly binds to *S. flexneri* 3b O-Ag (Figure 6C and Figure S6B). The variants showed the same pattern in relative apparent affinities as against *S. flexneri* 3a. Specifically, single amino acid variants S30G, L33N, and A51R all increased apparent affinity. N90H bound similarly to the parent antibody and H34A had a lower apparent affinity than the parent antibody. All combinations of the beneficial single mutations were found to be synergistic except for L33N + A51R. The combination of all 5 mutations bound poorly, similarly to H34A alone. The best binding single mutation was L33N, and the best binding combinatorial mutation was S30G + L33N, which had EC_50_ values of 322.5 nM and 165.3 nM respectively. We next wanted to determine whether these improvements in affinity for *S. flexneri* 3b O-Ag are sufficient to drive effector functions against this serotype. We compared the parent mAb to the best binding single variant, L33N, and the best binding combinatorial variant, S30G + L33N, in an L-ABA assay. In this assay, hFlex3a2_v2 displayed weak complement-mediated bactericidal activity against serotype 3b, only reaching 37% killing at 90 nM, the highest concentration of antibody tested (Figure 6D). Both the single L33N mutation and combination of S30G + L33N mutations were more potent in this assay, with IC_50_ values of 30.4 nM and 18.0 nM, respectively. Similarly, in an OPA assay, hFlex3a2_v2 was able to facilitate opsonophagocytosis at a concentration of 30 nM, leading to 7.4% CTV+ monocytes (Figure 6E). At the same concentration of mAb, the most potent single amino acid variants were ∼3-fold more potent at OPA, while the most potent combinatorial variants were ∼4.5-fold more potent (Figure 6E). However, despite improving affinity and potency in effector function assays, the higher affinity variants of hFlex3a2_v2 did not block *S. flexneri* 3b invasion (Figure 6F). Together, this data shows that panning against *S. flexneri* 3a was able to increase the functional breadth of this antibody to also exhibit effector functions against *S. flexneri* 3b, likely by improving the affinity to a shared epitope on these two serotypes that was weakly recognized by the parent mAb.

## Discussion

Here, we report the development of a panning strategy that enabled the rapid affinity maturation of a mAb against the O-Ag of *S. flexneri* 3a. We demonstrate that variants with improved affinity exhibit more potent effector functions against both *S. flexneri* 3a and the related but distinct serotype *S. flexneri* 3b. By pairing whole-cell in-solution panning with NGS and using both the target bacterial strain and a related strain with a completely unrelated O-Ag, we were able to efficiently identify specific mutations that could increase affinity against the target strain. It is notable that out of the four tested variants that met the criteria for specific enrichment (S30G, L33N, A51R, and N90H), three of them (S30G, L33N, A51R) improved affinity by ELISA, and all four exhibited improved potency in at least one effector function assay. One of these mutations, L33N, was sufficient to increase apparent affinity in ELISA by approximately 10-fold over the parent mAb, which is comparable to what has been reported for germline mAbs against *Chlamydia* core lipopolysaccharide (28). In contrast, H34A, which was enriched against both bacterial species (possibly indicating that this mutation improves a non-specific parameter such as display efficiency on phage), had lower affinity than the parent, supporting the rationale of this approach.

Unexpectedly, while variants in the hFlex3a2 VL library were successfully enriched following panning, no variants were enriched from the VH library (Figure S1). It is possible that the VH is already optimized, and we were simply unable to find point mutations that conferred additional advantage. This is supported by the observation that, while the VL protein sequence of hFlex3a2 is identical to germline, the VH sequence already contains somatic mutations that result in four amino acid substitutions (Figure 1A). However, the fact that no enrichment was observed against either *S. flexneri* 3a or *S. sonnei* (which would not be expected, as at least some variants should be nonspecifically enriched for reasons of improved display efficiency or scFv stability), leaves open the possibility that this was a technical issue with the panning process rather than a biological one. Further experiments would be needed to definitively determine whether any additional VH mutations can further improve binding and function of hFlex3a2. The identification and characterization of VH mutations that further synergize with the already identified VL mutations will be of interest in future studies.

As expected, higher affinity hFlex3a2 variants were generally more potent across all assays (Figure 5), with some notable exceptions. In L-ABA, affinity was tightly correlated with potency (Figure S5B), suggesting that potency in this assay has a linear relationship with affinity. Similarly, higher affinity variants were able to block invasion by *S. flexneri* 3a, while the parent mAb was not (Figure 5C). However, even these enhanced variants were unable to block invasion by serotype 3b (Figure 6F). While it is possible that the mechanism of action by which hFlex3a2 and its variants are inhibiting *S. flexneri* 3a invasion is not applicable to *S. flexneri* 3b, we suspect that this observation is more likely due to invasion inhibition requiring a greater affinity than L-ABA or OPA activity. Even though some hFlex3a2_v2 variants are more potent against *S. flexneri* 3b than the parental mAb, the best variants have weaker affinity and potency against serotype 3b than 3a. Given the inherent biological variability of the invasion assay compared to L-ABA or OPA, and that the parent mAb does not measurably inhibit invasion against even 3a, we think that the best variants simply lack sufficient affinity to reach the threshold where a decrease in invasion by 3b is consistently measurable in this assay. It is possible that panning against 3b with the hFlex3a2_v2 library would identify variants that further improve activity against this serotype.

The relationship between OPA and affinity was more complex than those between affinity and the L-ABA or invasion assay. While the highest affinity variants tested here (L33N and S30G + L33N) did exhibit higher degrees of opsonophagocytosis than the parent mAb, N90H, which does not have detectably higher affinity and was not more potent in L-ABA, also improved macrophage uptake of both *S. flexneri* 3a and 3b (Figure 5B and 6E). Similarly, H34A, which has lower affinity and L-ABA potency than the parent, performed similarly to the parent mAb in the OPA (Figure 5B and 6E). A similar phenomenon has been observed for some mAbs targeting the human immunodeficiency virus (HIV) glycoprotein gp120 as they exhibit different capacities to drive antibody-dependent cellular cytotoxicity against HIV-infected cells displaying gp120, despite being from the same lineage, having the same affinity, and binding the same epitope (44). In that study, it was proposed that these differences are a consequence of either subtle variations in the structural recognition of the epitope or other changes in the binding dynamics between the mAb and the antigen. We suspect similar factors, such as changes in the kinetics of binding (but not necessarily equilibrium affinity) may allow N90H to coordinate opsonophagocytosis more efficiently than the parent mAb despite having the same apparent equilibrium affinity. Unfortunately, because purified *S. flexneri* 3a O-Ag is present at a range of molecular weights, it is not currently possible to obtain accurate kinetic measurements to test this hypothesis. However, future studies on the relationship between affinity and effector function potency among mAbs against bacterial polysaccharides may provide more mechanistic insights. Altogether, these data support the hypothesis that higher affinity generally improves antibacterial function performance for anti-O-Ag mAbs, with some notable exceptions and potential caveats.

Most importantly, this work informs the development of therapeutic mAbs targeting bacterial polysaccharides. More test cases are required to establish the generalizability of this approach. However, these data demonstrate that in principle, this whole-cell in-solution panning approach can rapidly identify variants with improved affinity without the need for antigen purification, and these improvements lead to direct functional improvements. More interestingly, the observed enhancement of activity against *S. flexneri* 3b suggests that if a mAb is targeting an epitope shared with other bacterial serotypes, it may be possible to improve functional breadth alongside potency. This supports the development of clinically relevant mAbs against bacterial polysaccharides, because discovery campaigns can now prioritize mAbs that have relatively high breadth by binding sugars common to many bacterial polysaccharides. Broad binders can then be rapidly improved using this method. Future work will explore the potential of more complex panning strategies that incorporate multiple serotypes, which can further facilitate the identification of variants with improved breadth. Altogether, these results are an important step towards expanding the mAb engineering toolkit for anti-polysaccharide mAbs, which in turn may help develop novel therapeutics against AMR bacterial pathogens.

## Materials and methods

### Bacterial strains and growth conditions

Bacterial strains and plasmids used in this study can be found in Table S2. *Shigella* strains were maintained at −80 °C in tryptic soy broth (TSB) containing 20% (vol/vol) glycerol. *Shigella* was grown aerobically at 37 °C on TSB agar with 0.01% Congo red dye.

For all assays, bacteria were grown in the desired medium overnight, then subcultured 1:100 into fresh growth medium and grown aerobically at 250 rpm, 37 °C.

### Cell culture conditions

HeLa cells were grown in high glucose DMEM (Gibco) + 10% FBS + 2 mM glutamine and 1x Pen/Strep. THP-1 cells were grown in RPMI (Gibco) + 10% FBS + 10mM Hepes pH 7.4 + 1 mM Sodium Pyruvate + 55 µM beta-mercaptoethanol and 1x Pen/Strep. For assays involving bacteria, HeLa and THP-1 cells were first washed into fresh media that does not contain Pen/Strep. Expi293™ cells were grown in Expi 293™ Expression Medium (Gibco).

### 10x sequencing of hybridomas

An unsequenced hybridoma expressing the mAb hFlex3a2, which was discovered and described previously by WRAIR and collaborators (37), was provided to IAVI by WRAIR. Single cells were isolated using a BD FACSMelody™ cell sorter, and heavy and light chain variable domains were sequenced using the 10X single cell genomics platform (10X Genomics, Inc.). Sequence analysis to predict CDRs, align to germline, and identify rare residues was done using IMGT/V-Quest, IgBLAST, and abYsis (45–47). The VH and VL sequences were cloned into a modified pcDNA3.2 expression vector and purified as full IgG as described below. The mAb was confirmed to have properties consistent with the parental hybridoma mAb (Figure 1).

### Construction of plasmids

Primers and plasmids used in this study can be found in Table S3. For periplasmic expression of scFvs, hFlex3a2 and hFlex3a2_v2 were designed as scFvs in the following orientation — PelB signal sequence, hFlex3a2 VH, (GGGGS)3X linker, hFlex3a2 VL — ordered as gBlocks (Integrated DNA Technologies), then cloned between the NcoI and BamHI restriction sites of the bacterial expression vector pET28b (+) using NEBuilder® HiFi DNA Assembly following the manufacturer’s instructions (New England Biolabs). For expression as huIgG1, the variable regions of hFlex3a2 and hFlex3a2_v2 were cloned into a pcDNA3.2-based human antibody expression vectors containing the relevant huIgG1 or Ig kappa constant domains. For single amino acid mAb variants, primers were designed to insert the mutations into the appropriate VL sequence by HiFi DNA Assembly (New England BioLabs). Combinatorial variants of hFlex3a2 and hFlex3a2_v2 VL chains were ordered as gBlocks and inserted into the appropriate human antibody expression vector using NEBuilder® HiFi assembly.

### Construction of *Shigella* mutants

*Shigella flexneri* 3a and 3b *tolR* mutants that hyperproduce OMVs (48), and *S. flexneri* 3a *waaL* mutant (O-Ag knockout) (49) were produced by the gene-gorging method as previously described (50). Briefly, a mutation cassette consisting of 500 bp of *Shigella* genomic DNA upstream of the target mutation site, the kanamycin resistance gene from plasmid pKD4, and 500 bp of DNA downstream of the target mutation site, flanked on both sides of the mutation cassette by the restriction site I-SceI was constructed using splice by overlap extension PCR (51). The mutation cassette was A-tailed and ligated into the plasmid pGEM®-T Easy (Promega) to create a donor plasmid. The donor plasmid was electroporated into the desired *S. flexneri* strain that had previously been electroporated with plasmid pACBSR. Strains containing both the donor plasmid and pACBSR were grown for 6 hours at 30 °C in LB + chloramphenicol + 0.4% arabinose to induce genomic recombination and then plated on TSBA + Kanamycin for single colonies. Colonies were passaged multiple times at 37 °C to promote the loss of pACBSR. After multiple passages, colonies were confirmed to be kanamycin resistant but chloramphenicol sensitive, indicating loss of donor plasmid, and the successful insertion of the genomic mutation was verified by Sanger sequencing.

### Phage library construction and panning

Double-barcoded site-saturated mutagenesis scanning libraries of the VL and VH of hFlex3a2_v2 were designed and ordered from Twist Biosciences. The libraries were amplified with primers to introduce SfiI sites and then ligated into the SfiI site of phagemid pCOMB3XSS (52). To produce phage, the phagemid libraries were electroporated into XL1-Blue (Agilent), recovered in 5 mL of SOC, and then grown (37 °C, 250 RPM) to OD_600_ ∼0.5 in 10 mL of Super Broth + Tet + Carb before being super infected with M13ko7 helper phage (New England Biosciences) at an MOI of 20, expanded, and grown overnight (30°C, 250 rpm) in 200 mL of super broth + Tet + Carb + Kan.

Phage was precipitated from the supernatant with 0.5 M NaCl and 4% PEG-8,000 as previously described (53) and then panned against whole bacteria in-solution, based on a previously published method (42). Briefly, 1 x 10^9^ cfu of *S. flexneri* 3a or *S. sonnei* were blocked in 3% nonfat milk in PBS and then each bacterial species was incubated with 5 x 10^10^ pfu of blocked phage particles for 15 min to 2 h. Non-binding phage was washed 5x with PBS + 5% Tween-20 after round 1 of panning and washed 10x after round 2. Binding phage was eluted by mixing with 0.1 M HCl pH 2.2 (pH adjusted with glycine) for ten minutes at room temperature, then neutralized with 2 M Tris base and used to reinfect XL1-Blue cells to generate phage for the second round of panning. XL1-Blue containing the input library, post-panning round 1 phagemids, and post-panning round 2 phagemids were maxiprepped, the variable chain sequences were PCR-amplified and deep-sequenced by MiSeq (Illumina) using Amplicon-EZ platform (Genewiz). The relative enrichment of phage library variants against *S. flexneri* 3A and *S. sonnei* after rounds 1 and 2 of panning was quantified using an in-house Python script.

### Antibody and antigen purification

For purification of IgGs, Expi293 cells were co-transfected with the desired heavy chain and light chain pair at a ratio of 1:2.5 with FectoPRO® (Polyplus) in Expi293^TM^ Expression Medium (Gibco). The cells were supplemented with 3 mM valproic acid (Sigma-Aldrich) and 0.4% D-(+)-glucose (Sigma-Aldrich) 24 h after transfection. The supernatant was harvested on day 5 post-transfection, IgGs were purified using Praesto AP+ agarose resin (Purolite), and then buffer exchanged and concentrated using 30 kDa cutoff spin filter (Amicon).

scFv purification was based on a previously described method (54). Briefly, BL21 containing the desired scFv cloned into pET28b (+) was grown to OD_600_ 0.6-0.8 in terrific broth (37 °C, 250 rpm) and then induced with 1 mM IPTG overnight (18 °C, 250 RPM). Bacteria were collected, resuspended in 40 mL of TES buffer (0.5 M Sucrose, 0.5 mM EDTA, and 0.2 M Tris-HCl pH 8.0), incubated at 4 °C for 30 min, and then osmotically shocked with 80 mL of ice-cold ddH_2_O for 1 h at 4 °C. Lysed bacteria were pelleted, the supernatant was collected, adjusted to 150 mM NaCl, 2 mM MgCl_2_, 20 mM imidizaole, and incubated with 1 mL of Ni-NTA resin (Anatrace) at 4 °C for at least 1.5 h. scFv bound resin was packed onto gravity columns, washed extensively with wash buffer (150 mM NaCl, 2 mM MgCl_2_, 30 mM imidazole), and then eluted with elution buffer (150 mM NaCl, 2 mM MgCl_2_, 300 mM imidazole) before buffer exchanging and concentrating with PBS using a 10 kDa cutoff centrifugal filter (Amicon).

To purify OMVs, supernatant was collected from *S. flexneri* 3a or 3b tolR::kan mutants grown overnight in SSDM (48) supplemented with 40 mg/L l-methionine and 20 mg/L tryptophan. The supernatant was filter-sterilized through a 0.2 µM filter (Sartorius) and then concentrated to ∼ 50 mL by tangential flow filtration using a Vivaflow® 100 kDa cutoff filter (Sartorius^®^). The concentrated supernatant was ultracentrifuged at 200,000 x g for 1.5 h to pellet OMVs, which were then resuspended in 1 mL of PBS and stored at −20 °C for future use. Total protein content of OMVs was quantified by Micro-BCA^TM^ assay (Thermo Fisher) following the manufacturer’s instructions. OMV amount was normalized by total protein content for ELISA assays.

O-Ag was purified from whole bacteria by acetic acid hydrolysis (55). Acetic acid was added at a final concentration of 2% to a culture of stationary phase *S. flexneri* 3a and incubated at 100 °C for 5 h. After hydrolysis, the culture was neutralized to pH ∼ 6 with ammonium hydroxide (Sigma-Aldrich). Bacterial debris was pelleted, the supernatant was buffer exchanged by tangential flow filtration using a VivaFlow 30 kDa cutoff filter (Sartorius) first into 1 M NaCl, and then into ddH_2_O.The buffer exchanged supernatant was concentrated to ∼50 mL. Citrate buffer (20 mM, pH 3) was added to the supernatant, and it was incubated at room temperature for 30 min, centrifuged at 12,000 x g for 30 min and then passed through a Sartobind® IEX S75 cationic exchange filter (Sartorius). After cation exchange, the supernatant was precipitated once more with 18 mM Na_2_HPO_4_, 24% ethanol, and 200 mM CaCl_2_ at room temperature for 30 min, then buffer exchanged as described above before being concentrated to approximately 1 mL in ddH_2_O using a 30 kDa cutoff spin filter (Amicon). The purified O-Ag was quantified using a Total Carbohydrate Assay Kit (Sigma-Aldrich) following the manufacturer’s instructions. Contaminating DNA and protein levels were assessed by nanodrop analysis and Micro-BCA™ assay, respectively, and confirmed to be at least 10-fold lower than carbohydrate concentration.

### ELISA assays

All ELISA assays were performed in 384-well high binding ELISA plates (Corning). Wells were coated overnight at 4 °C with antigen in PBS at a concentration of 2.5 µg/mL for OMVs or 5 µg/mL for ovalbumin, CHO-SMP, insulin, and ssDNA. Antigen-coated wells were blocked in PBS containing 3% BSA and 0.05% Tween-20 at 37 °C for 1 h. Antibodies to be tested were diluted in 1% BSA + 0.017% Tween-20 at the desired antibody concentration and incubated with the blocked antigen at 37 °C for 1 h. After incubation with the test antibody, wells were washed 3x with PBS + 0.05% Tween-20 and incubated with the appropriate secondary antibody at room temperature for 1 h. Secondary antibodies used were either AP-conjugated anti-Flag M2 (Sigma-Aldrich) for scFvs or AP-conjugated anti-Human Fc (Jackson ImmunoResearch) for huIgG1s, both at a 1:1000 dilution in PBS + 1% BSA + 0.017% Tween-20. After washing three times more, AP substrate tablets (Sigma-Adrich) were dissolved and diluted to 1 mg/mL in AP buffer (50 mM MgCl_2_, 100 mM NaCl, 1%Tween 20). ELISAs were developed with AP substrate (20 min for IgGs or 40 min for scFvs) at room temperature. Absorbance at 405 nm was read on a BioTek Synergy microplate reader.

### Luminescent Antibody Bactericidal Assays (L-ABA)

L-ABA was adapted from previously published protocols (37,56). Briefly, the desired *S. flexneri* serotype was grown to mid-log phase in TSB (BD Biosciences) and resuspended at a concentration of 3 x 10^5^ CFU/mL in 100 µL of assay buffer (HBSS + 0.1% Gelatin, pH 7.4) with 7.5% Baby Rabbit Complement (Pel-Freez Biologicals), and the test antibody at the desired concentration. The L-ABA reaction was incubated for 3 h at 37 °C, washed once with 100 μL of fresh assay buffer, mixed 1:1 with BacTiter-Glo™ (Promega), and incubated for 5 min at room temperature. Luminescence was read on a BioTek Synergy plate reader. Adalimumab (57) expressed as a chimeric macaque IgG1 was used as a control mAb. To normalize for differences in bacterial growth, the data was fit with a sigmoidal four parameter logistic curve on GraphPad Prism. Each curve was normalized to the Top best-fit value of their curve to determine their fraction maximal luminescence. Control mAb values are normalized to the luminescence signal of total bacterial growth in assay buffer and baby rabbit complement.

### Opsonophagocytosis Assays (OPA)

For OPA, the desired *S. flexneri* strain was grown to OD_600_ of approximately 0.5 in TSB, washed with PBS, and stained with a 1:50 dilution of CellTrace™ Violet (CTV, Thermo Fisher) for 15 min at 37 °C. Excess dye was quenched with FACS buffer (PBS + 1% FBS), the bacteria were washed in PBS and resuspended in assay buffer (HBSS + 0.1% gelatin, pH 7.4). 2.5 x 10^6^ CFU of bacteria were mixed with the desired concentration of mAb in assay buffer and incubated at 37 °C for 30 min. Opsonized bacteria were then mixed with 5 x 10^4^ THP-1 cells and incubated for 30 min at 37 °C in 5% CO_2_. After 30 min, the OPA reactions were stained on ice in the dark for 30 min with Live/Dead Near-IR dye (Thermo Fisher) diluted 1:2000 in PBS and FITC-conjugated anti-CD45 antibody (BD Biosciences) diluted 1:4000. The cells were then fixed with 4% PFA for 30 min, on ice. To determine relative OPA activity, the percentage of THP-1 cells incubated with *S. flexneri* opsonized with the mAbs of interest that were Near IR-, CD45+, and CTV+ was quantified by flow cytometry using a FACSCelesta (BD Biosciences).

### Invasion Assays

Invasion was measured by a gentamicin protection assay (58). *S. flexneri* was grown to an OD_600_ of approximately 0.7 in TSB (37 °C, 250 RPM), washed in PBS, resuspended at a concentration of 1 x 10^9^ CFU/mL, and incubated with 1 µM of the desired mAb or PBS (control) for 30 min at 37 °C. 10 µL of bacteria (∼1 x 10^7^ CFU, MOI = 20) was added to a monolayer of approximately 5 x 10^5^ HeLa cells in a tissue culture-treated 24-well plate (Corning) and centrifuged at 900 x g for 10 min. The plate was incubated for 30 min at 37 °C and 5% CO_2_, washed two times with PBS, the media was replaced with DMEM containing 25 µg/mL gentamicin, and the plate was incubated for an additional 1 h. The wells were washed twice more with PBS and then lysed with 0.1% Triton X-100. Serial dilutions of the mAb-treated and control bacteria were plated, and percent invasion was calculated. Adalimumab (57) expressed as a chimeric macaque IgG1 was used as a control mAb.

### Bacterial surface staining

To measure bacterial surface staining, *S. flexneri* was grown to an OD_600_ of approximately 0.5, washed and resuspended in PBS to a concentration of 2 x 10^9^ CFU/mL. The desired mAb was added to 25 µL of bacteria at a final concentration of 50 nM and incubated at 37 °C for 30 min. The bacteria were washed with PBS and then incubated with a 1:100 dilution of the appropriate secondary antibody for 30 min at room temperature. For scFvs, the secondary antibody was Alexa Fluor 647-conjugated anti-Flag antibody (Abcam), and for huIgGs this was BV421-conjugated anti-Human IgG (BD Biosciences). Bacteria were then fixed with 4% PFA for 30 min on ice. Median fluorescence intensity was determined by flow cytometry on a FACSSymphony A5 (BD Biosciences).

### Biolayer interferometry

Binding of hFLex3a2_v2 variants to *S. flexneri* 3a O-Ag was characterized by BLI using an Octet® R8 (Sartorius) with Octet AHC Biosensors (Sartorius). All antibodies and antigens were in PBS + 0.02% Tween-20 + 0.01% BSA. The BLI protocol was as follows: 300 seconds baseline, 120 seconds loading with 1 μM mAb, 180 seconds baseline, 360 seconds association with 100 μg/mL of *S. flexneri* 3a O-Ag or 540 seconds with 20 μg/mL of *S. flexnei* 3a O-Ag, and 600 seconds dissociation. BLI results were analyzed on Octet Analysis Studio (Sartorius) and plotted with GraphPad Prism. hFlex2a1 (37) expressed as a chimeric huIgG1 was used as a control mAb.

### O-Ag visualization

To visualize O-Ag, crude protein-free LPS extract was purified from *S. flexneri* 3a and *S. flexneri* 3a waaL::kan (O-Ag-) following the method of Darveau and Hancock (59). Bacterial LPS extract was separated by SDS-PAGE and stained with Pro-Q™ Emerald 300 Lipopolysaccharide Gel Stain Kit (Thermo Fisher) following the manufacturer’s instructions. Stained gels were visualized on a ChemiDoc MP Imaging System (Bio-Rad).

---

**Fig S1**: **Mutation enrichment in the VH library after panning.** (A) Fold-change in the frequency of next-generation sequencing reads for variant mutations at each VH site of hFlex3a2_v2 after two rounds of panning, relative to their frequency in the input library. (B) log10 enrichment of variants in the VH library after two rounds of panning against *S. flexneri* 3a and *S. sonnei*. Dotted line is 20 standard deviations above the mean mutation enrichment in the input library and indicates a stringent threshold set to distinguish bacteria-mediated enrichment from background variability.

**Fig S2: Distribution of next-generation sequencing reads across the VL library variants after panning.** A heatmap showing the percentage of all reads belonging to the indicated hFlex3a2_v2 variant after two rounds of panning against *S. flexneri* 3a.

**Fig S3: Enrichment of VL mutations for increased affinity to the surface of *S. flexneri 3a* after two rounds of panning**. Enrichment of the indicated amino acids in the library variants after panning against *S. flexneri* 3a was calculated relative to the enrichment against *S. sonnei*. All data is log10 transformed. White squares with an “X” through them indicate variants for which no reads were detected in either the input or output libraries, suggesting that these variants may not have been successfully synthesized.

**Fig S4: hFlex3a2_v2 variant *S. flexneri* 3a OMV and polyreactiviy ELISAs** (A) The same ELISA as Figure 4B is reproduced here, but with all variants labeled. (B) ELISA measuring non-specific binding of hFlex3a2_v2 variants to 5 μg/mL of insulin, CHO-SMPs, or ssDNA. A known poly-reactive mAb (4E10 targeting HIV-1) and a known non-polyreactive mAb (Adalimumab) serve as positive and negative controls, respectively. This experiment is the result of three replicates. (C) Gating strategy for quantifying OPA. Cells are gated first by size, then by CD45+ Dead-populations. An unrelated, nonbinding mAb negative control is used to determine the %CTV+ gate.

**Fig S5: L-ABA analysis of hFlex3a2_v2 variants against *S. flexneri* 3a** (A) L-ABA from Fig 5A is reproduced here, but with all variants labeled. (B) log10 IC_50_ from L-ABA and EC_50_ from ELISA are correlated. R^2^ represents Pearson correlation coefficient.

**Fig S6: hFlex3a2_v2 variant ELISA against *S. flexneri* 3b OMVs.** (A) SDS-PAGE of crude LPS extract from *S. flexneri* 3a and *S. flexneri* 3a waaL::kan (O-Ag-) confirms that knocking out waaL eliminates O-Ag expression. Gel was stained with Pro-Q Emerald 3000. (B)The ELISA from Fig 6C is reproduced here, but with all variants labeled.

## Acknowledgements

The hFlex3a2 mAb was generated by the laboratory of Dr. Moon Nahm at the University of Alabama Birmingham in collaboration with the Walter Reed Army Institute of Research (36). We thank collaborators from the Wisconsin National Primate Research Center, Jennifer Hayes, Dr. Faye Hartmann, and Dr. Eric Peterson, for sharing isolates WS-016 and WS-037. We thank Finora Franck for invaluable program management support, Dr. Jon Heinrichs and Dr. Tawanda Mandizvo for a critical reading of this manuscript, Katherine McKenney for assistance with 10X sequencing, and Mark Ochoa for assistance with cell culture. We also thank Dr. Stephen Baker and Dr. Timothy Scott for scientific advice and support, as well as providing the Shigella sonnei 53G lvp::kan strain. Finally, we would like to thank current and former members of the RESISTANT Research Steering Group (including Wellcome Trust Program Officers) for their invaluable support of this work, including Dr. Pete Gardner, Dr. Colleen Loynachan, Dr. Kate Mills, Dr. Nathalie Herze-Vourc’h, and Dr. Christiane Gerke. This work was supported by the Wellcome Trust RESISTANT grant.

## Works Cited

1. GBD 2021 Antimicrobial Resistance Collaborators. Global burden of bacterial antimicrobial resistance 1990-2021: a systematic analysis with forecasts to 2050. Lancet Lond Engl. 2024 Sept 28;404(10459):1199–226.

2. La Guidara C, Adamo R, Sala C, Micoli F. Vaccines and Monoclonal Antibodies as Alternative Strategies to Antibiotics to Fight Antimicrobial Resistance. Int J Mol Sci. 2024 May 17;25(10):5487.

3. Perrett KP, Nolan TM, McVernon J. A Licensed Combined Haemophilus influenzae Type b-Serogroups C and Y Meningococcal Conjugate Vaccine. Infect Dis Ther. 2013 June;2(1):1– 13.

4. Ladhani SN. Two decades of experience with the Haemophilus influenzae serotype b conjugate vaccine in the United Kingdom. Clin Ther. 2012 Feb;34(2):385–99.

5. Hedari CP, Khinkarly RW, Dbaibo GS. Meningococcal serogroups A, C, W-135, and Y tetanus toxoid conjugate vaccine: a new conjugate vaccine against invasive meningococcal disease. Infect Drug Resist. 2014;7:85–99.

6. Grijalva CG, Nuorti JP, Arbogast PG, Martin SW, Edwards KM, Griffin MR. Decline in pneumonia admissions after routine childhood immunisation with pneumococcal conjugate vaccine in the USA: a time-series analysis. Lancet Lond Engl. 2007 Apr 7;369(9568):1179– 86.

7. Milligan R, Paul M, Richardson M, Neuberger A. Vaccines for preventing typhoid fever. Cochrane Database Syst Rev. 2018 May 31;5(5):CD001261.

8. Baker S, Krishna A, Higham S, Naydenova P, O’Leary S, Scott JB, et al. Exploiting human immune repertoire transgenic mice for protective monoclonal antibodies against antimicrobial resistant Acinetobacter baumannii. Nat Commun. 2024 Sept 12;15(1):7979.

9. Aguiar-Alves F, Le HN, Tran VG, Gras E, Vu TTT, Dong OX, et al. Antivirulence Bispecific Monoclonal Antibody-Mediated Protection against Pseudomonas aeruginosa Ventilator-Associated Pneumonia in a Rabbit Model. Antimicrob Agents Chemother. 2022 Feb 15;66(2):e0202221.

10. Nielsen TB, Yan J, Slarve M, Lu P, Li R, Ruiz J, et al. Monoclonal Antibody Therapy against Acinetobacter baumannii. Infect Immun. 2021 Sept 16;89(10):e0016221.

11. Nielsen TB, Yan J, Luna BM, Talyansky Y, Slarve M, Bonomo RA, et al. Monoclonal Antibody Requires Immunomodulation for Efficacy Against Acinetobacter baumannii Infection. J Infect Dis. 2021 Dec 15;224(12):2133–47.

12. Richards AF, Doering JE, Lozito SA, Varrone JJ, Willsey GG, Pauly M, et al. Inhibition of invasive salmonella by orally administered IgA and IgG monoclonal antibodies. PLoS Negl Trop Dis. 2020 Mar;14(3):e0007803.

13. Gulati S, Beurskens FJ, de Kreuk BJ, Roza M, Zheng B, DeOliveira RB, et al. Complement alone drives efficacy of a chimeric antigonococcal monoclonal antibody. PLoS Biol. 2019 June;17(6):e3000323.

14. Kalfopoulou E, Laverde D, Miklic K, Romero-Saavedra F, Malic S, Carboni F, et al. Development of Opsonic Mouse Monoclonal Antibodies against Multidrug-Resistant Enterococci. Infect Immun. 2019 Sept;87(9):e00276–19.

15. Pennini ME, De Marco A, Pelletier M, Bonnell J, Cvitkovic R, Beltramello M, et al. Immune stealth-driven O2 serotype prevalence and potential for therapeutic antibodies against multidrug resistant Klebsiella pneumoniae. Nat Commun. 2017 Dec 8;8(1):1991.

16. Rollenske T, Szijarto V, Lukasiewicz J, Guachalla LM, Stojkovic K, Hartl K, et al. Cross-specificity of protective human antibodies against Klebsiella pneumoniae LPS O-antigen. Nat Immunol. 2018 June;19(6):617–24.

17. Parzych EM, Gulati S, Zheng B, Bah MA, Elliott STC, Chu JD, et al. Synthetic DNA Delivery of an Optimized and Engineered Monoclonal Antibody Provides Rapid and Prolonged Protection against Experimental Gonococcal Infection. mBio. 2021 Mar 16;12(2):e00242–21.

18. Que YA, Lazar H, Wolff M, François B, Laterre PF, Mercier E, et al. Assessment of panobacumab as adjunctive immunotherapy for the treatment of nosocomial Pseudomonas aeruginosa pneumonia. Eur J Clin Microbiol Infect Dis Off Publ Eur Soc Clin Microbiol. 2014 Oct;33(10):1861–7.

19. Chastre J, François B, Bourgeois M, Komnos A, Ferrer R, Rahav G, et al. Safety, efficacy, and pharmacokinetics of gremubamab (MEDI3902), an anti-Pseudomonas aeruginosa bispecific human monoclonal antibody, in P. aeruginosa-colonised, mechanically ventilated intensive care unit patients: a randomised controlled trial. Crit Care Lond Engl. 2022 Nov 15;26(1):355.

20. Healthcare G. ATS 2025: promising results with gremubamab in bronchiectasis [Internet]. Clinical Trials Arena. 2025 [cited 2025 Sept 8]. Available from: https://www.clinicaltrialsarena.com/analyst-comment/ats-2025-promising-results-gremubamab-bronchiectasis/

21. Erridge C, Bennett-Guerrero E, Poxton IR. Structure and function of lipopolysaccharides. Microbes Infect. 2002 July;4(8):837–51.

22. Formal SB, Oaks EV, Olsen RE, Wingfield-Eggleston M, Snoy PJ, Cogan JP. Effect of prior infection with virulent Shigella flexneri 2a on the resistance of monkeys to subsequent infection with Shigella sonnei. J Infect Dis. 1991 Sept;164(3):533–7.

23. Cohen D, Meron-Sudai S, Bialik A, Asato V, Goren S, Ariel-Cohen O, et al. Serum IgG antibodies to Shigella lipopolysaccharide antigens - a correlate of protection against shigellosis. Hum Vaccines Immunother. 2019;15(6):1401–8.

24. Kappler K, Hennet T. Emergence and significance of carbohydrate-specific antibodies. Genes Immun. 2020 Aug;21(4):224–39.

25. Haji-Ghassemi O, Blackler RJ, Martin Young N, Evans SV. Antibody recognition of carbohydrate epitopes. Glycobiology. 2015 Sept;25(9):920–52.

26. Borenstein-Katz A, Warszawski S, Amon R, Eilon M, Cohen-Dvashi H, Leviatan Ben-Arye S, et al. Biomolecular Recognition of the Glycan Neoantigen CA19-9 by Distinct Antibodies. J Mol Biol. 2021 July 23;433(15):167099.

27. Amon R, Rosenfeld R, Perlmutter S, Grant OC, Yehuda S, Borenstein-Katz A, et al. Directed Evolution of Therapeutic Antibodies Targeting Glycosylation in Cancer. Cancers. 2020 Sept 30;12(10):2824.

28. Haji-Ghassemi O, Müller-Loennies S, Brooks CL, MacKenzie CR, Caveney N, Van Petegem F, et al. Subtle Changes in the Combining Site of the Chlamydiaceae-Specific mAb S25-23 Increase the Antibody-Carbohydrate Binding Affinity by an Order of Magnitude. Biochemistry. 2019 Feb 12;58(6):714–26.

29. Hu J, Huang X, Ling CC, Bundle DR, Cheung NKV. Reducing epitope spread during affinity maturation of an anti-ganglioside GD2 antibody. J Immunol Baltim Md 1950. 2009 Nov 1;183(9):5748–55.

30. Thomas R, Patenaude SI, MacKenzie CR, To R, Hirama T, Young NM, et al. Structure of an anti-blood group A Fv and improvement of its binding affinity without loss of specificity. J Biol Chem. 2002 Jan 18;277(3):2059–64.

31. Brorson K, Thompson C, Wei G, Krasnokutsky M, Stein KE. Mutational analysis of avidity and fine specificity of anti-levan antibodies. J Immunol Baltim Md 1950. 1999 Dec 15;163(12):6694–701.

32. Deng SJ, MacKenzie CR, Sadowska J, Michniewicz J, Young NM, Bundle DR, et al. Selection of antibody single-chain variable fragments with improved carbohydrate binding by phage display. J Biol Chem. 1994 Apr 1;269(13):9533–8.

33. Brummell DA, Sharma VP, Anand NN, Bilous D, Dubuc G, Michniewicz J, et al. Probing the combining site of an anti-carbohydrate antibody by saturation-mutagenesis: role of the heavy-chain CDR3 residues. Biochemistry. 1993 Feb 2;32(4):1180–7.

34. Yelton DE, Rosok MJ, Cruz G, Cosand WL, Bajorath J, Hellström I, et al. Affinity maturation of the BR96 anti-carcinoma antibody by codon-based mutagenesis. J Immunol Baltim Md 1950. 1995 Aug 15;155(4):1994–2004.

35. Wang W, Bunyatov M, Moffat D, Lopez-Barbosa N, DeLisa MP. Engineering affinity-matured variants of an anti-polysialic acid monoclonal antibody with superior cytotoxicity-mediating potency. Cell Chem Biol. 2025 Aug 21;32(8):1042–1057.e6.

36. Deng SJ, MacKenzie CR, Hirama T, Brousseau R, Lowary TL, Young NM, et al. Basis for selection of improved carbohydrate-binding single-chain antibodies from synthetic gene libraries. Proc Natl Acad Sci U S A. 1995 May 23;92(11):4992–6.

37. Lin J, Smith MA, Benjamin WH, Kaminski RW, Wenzel H, Nahm MH. Monoclonal Antibodies to Shigella Lipopolysaccharide Are Useful for Vaccine Production. Clin Vaccine Immunol CVI. 2016 Aug;23(8):681–8.

38. Mancini F, Gasperini G, Rossi O, Aruta MG, Raso MM, Alfini R, et al. Dissecting the contribution of O-Antigen and proteins to the immunogenicity of Shigella sonnei generalized modules for membrane antigens (GMMA). Sci Rep. 2021 Jan 13;11(1):906.

39. Kang TH, Seong BL. Solubility, Stability, and Avidity of Recombinant Antibody Fragments Expressed in Microorganisms. Front Microbiol. 2020;11:1927.

40. Maynard JA, Maassen CBM, Leppla SH, Brasky K, Patterson JL, Iverson BL, et al. Protection against anthrax toxin by recombinant antibody fragments correlates with antigen affinity. Nat Biotechnol. 2002 June;20(6):597–601.

41. Chowdhury PS, Vasmatzis G. Engineering scFvs for improved stability. Methods Mol Biol Clifton NJ. 2003;207:237–54.

42. Berry SK, Rust S, Caceres C, Irving L, Bartholdson Scott J, Tabor DE, et al. Phenotypic whole-cell screening identifies a protective carbohydrate epitope on Klebsiella pneumoniae. mAbs. 2022;14(1):2006123.

43. Perepelov AV, Shekht ME, Liu B, Shevelev SD, Ledov VA, Senchenkova SN, et al. Shigella flexneri O-antigens revisited: final elucidation of the O-acetylation profiles and a survey of the O-antigen structure diversity. FEMS Immunol Med Microbiol. 2012 Nov;66(2):201–10.

44. Doepker LE, Danon S, Harkins E, Ralph DK, Yaffe Z, Garrett ME, et al. Development of antibody-dependent cell cytotoxicity function in HIV-1 antibodies. eLife. 2021 Jan 11;10:e63444.

45. Swindells MB, Porter CT, Couch M, Hurst J, Abhinandan KR, Nielsen JH, et al. abYsis: Integrated Antibody Sequence and Structure-Management, Analysis, and Prediction. J Mol Biol. 2017 Feb 3;429(3):356–64.

46. Brochet X, Lefranc MP, Giudicelli V. IMGT/V-QUEST: the highly customized and integrated system for IG and TR standardized V-J and V-D-J sequence analysis. Nucleic Acids Res. 2008 July 1;36(Web Server issue):W503-508.

47. Ye J, Ma N, Madden TL, Ostell JM. IgBLAST: an immunoglobulin variable domain sequence analysis tool. Nucleic Acids Res. 2013 July;41(Web Server issue):W34-40.

48. Berlanda Scorza F, Colucci AM, Maggiore L, Sanzone S, Rossi O, Ferlenghi I, et al. High yield production process for Shigella outer membrane particles. PloS One. 2012;7(6):e35616.

49. Han W, Wu B, Li L, Zhao G, Woodward R, Pettit N, et al. Defining function of lipopolysaccharide O-antigen ligase WaaL using chemoenzymatically synthesized substrates. J Biol Chem. 2012 Feb 17;287(8):5357–65.

50. Herring CD, Glasner JD, Blattner FR. Gene replacement without selection: regulated suppression of amber mutations in Escherichia coli. Gene. 2003 June 5;311:153–63.

51. Heckman KL, Pease LR. Gene splicing and mutagenesis by PCR-driven overlap extension. Nat Protoc. 2007;2(4):924–32.

52. Andris-Widhopf J, Rader C, Steinberger P, Fuller R, Barbas CF. Methods for the generation of chicken monoclonal antibody fragments by phage display. J Immunol Methods. 2000 Aug 28;242(1–2):159–81.

53. Silva RP, DiVenere AM, Amengor D, Maynard JA. Antibodies binding diverse pertactin epitopes protect mice from Bordetella pertussis infection. J Biol Chem. 2022 Mar;298(3):101715.

54. Cai H, Yao H, Li T, Hutter CAJ, Li Y, Tang Y, et al. An improved fluorescent tag and its nanobodies for membrane protein expression, stability assay, and purification. Commun Biol. 2020 Dec 10;3(1):753.

55. Micoli F, Rondini S, Gavini M, Pisoni I, Lanzilao L, Colucci AM, et al. A scalable method for O-antigen purification applied to various Salmonella serovars. Anal Biochem. 2013 Mar 1;434(1):136–45.

56. Caradonna V, Pinto M, Alfini R, Giannelli C, Iturriza M, Micoli F, et al. High-Throughput Luminescence-Based Serum Bactericidal Assay Optimization and Characterization to Assess Human Sera Functionality Against Multiple Shigella flexneri Serotypes. Int J Mol Sci. 2024 Oct 16;25(20):11123.

57. Weinblatt ME, Keystone EC, Furst DE, Moreland LW, Weisman MH, Birbara CA, et al. Adalimumab, a fully human anti-tumor necrosis factor alpha monoclonal antibody, for the treatment of rheumatoid arthritis in patients taking concomitant methotrexate: the ARMADA trial. Arthritis Rheum. 2003 Jan;48(1):35–45.

58. Elsinghorst EA. Measurement of invasion by gentamicin resistance. Methods Enzymol. 1994;236:405–20.

59. Darveau RP, Hancock RE. Procedure for isolation of bacterial lipopolysaccharides from both smooth and rough Pseudomonas aeruginosa and Salmonella typhimurium strains. J Bacteriol. 1983 Aug;155(2):831–8.

