## Supplementary material for "Improvement of affinity and potency of a monoclonal antibody against *Shigella flexneri* 3a O-antigen via phage display and whole-cell in-solution panning": Table S1

| hFlex3a2 protein sequence | |
| --- | --- |
| Heavy Chain | QVQLQQSAPELARPGASVKMSCKASGYTFTSYTIHWVKQRPGQGLEWIGYISPSSGYTEYNQKFKDKTTLTADKSSITAYMQLSSLTSEDSAVYYCARLDNNYVYFDYWGLGTTLTVSS |
| Light Chain | DIVMTQSPATLSVTPGDRVSLSCRASQSISDYLHWYQQKSHESPRLLIKYASQSISGIPSRFSGSGSGSDFTLSINSVEPEDVGVYYCQNGHSFPLTFGAGTKLELK |
| hFlex3a2_v2 protein sequence | |
| Heavy Chain | QVQLQQS**G**PELARPGASVKMSCKASGYTFTSYTIHWVKQRPGQGLEWIGYISPSSGYTEYNQKFKDKTTLTADKSS**S**TAYMQLSSLTSEDSAVYYCARLDNNYVYFDYWGLGTTLTVSS |
| Light Chain | DIVMTQSPATLSVTPGDRVSLSCRASQSISDYLHWYQQKS**G**ESPRLLIKYASQSISGIPSRFSGSGSGSDFTLSINSVEPEDVGVYYCQNGHSFPLTFGAGTKLELK |

Underlined residues indicate sites that are different between hFlex3a2 and hFlex3a2_v2

Red residues indicate sites of amino acid changes for affinity matured variants:

S30G, L33N, H34A, A51R, and N90H
