## Supplementary material for "Improvement of affinity and potency of a monoclonal antibody against *Shigella flexneri* 3a O-antigen via phage display and whole-cell in-solution panning": Table S2

**Table S2.** Strains and plasmids used in this study.

| **Strain or plasmid** | **Description** | **Reference/Source** |
| --- | --- | --- |
| ***E. coli*** |  |  |
| NEB5α | Cloning strain | New England Biolabs |
| XL1-Blue | Phage library construction | Agilent Technologies |
| BL21 | Protein expression | New England Biolabs |
| ***Shigella*** |  |  |
| *S. flexneri* 210059 | Serotype 1b | (1) |
| *S. flexneri* 2457T | Serotype 2a | WRAIR |
| *S. flexneri* CDC 3591-52 | Serotype 2b | ATCC 12022 |
| *S. flexneri* J17B | Serotype 3a | WRAIR |
| *S. flexneri* WS-016 | Serotype 3b | This study |
| *S. flexneri* WS-037 | Serotype Y | This study |
| *S. sonnei* 53G lvp::kan | LVP stable *S. sonnei* | Stephen Baker Lab |
| *S. flexneri* J17B *tolR*::kan | tolR knockout, overproduces serotype 3a OMVs | This study |
| *S. flexneri* J17B *waaL*::kan | waaL knockout, serotype 3a O-Ag negative strain | This study |
| *S. flexneri* WS-037 *tolR*::kan | tolR knockout, overproduces serotype 3b OMVs | This study |
| **Human** |  |  |
| HeLa | Virulence assays | ATCC CCL-2 |
| Expi293 | Antibody transfection | Thermo Fisher Scientific |
| THP-1 | OPA | ATCC TIB-202 |
| **Plasmids** |  |  |
| pACBSR | Gene-gorging mutagenesis vector | (2) |
| pKD4 | Kan resistance casette | (3) |
| pGEMTeasy | Donor plasmid for gorging | Promega |
| pET28b (+) | Protein expression vector | Sigma Aldrich |
| pCOMB3XSS | Phage display vector | (4) |

Works Cited:

1.	Tansarli GS, Long DR, Waalkes A, Bourassa LA, Libby SJ, Penewit K, Almazan J, Matsumoto J, Bryson-Cahn C, Rietberg K, Dell BM, Hatley NV, Salipante SJ, Fang FC. 2023. Genomic reconstruction and directed interventions in a multidrug-resistant Shigellosis outbreak in Seattle, WA, USA: a genomic surveillance study. Lancet Infect Dis 23:740–750.

2.	Herring CD, Glasner JD, Blattner FR. 2003. Gene replacement without selection: regulated suppression of amber mutations in Escherichia coli. Gene 311:153–163.

3.	Datsenko KA, Wanner BL. 2000. One-step inactivation of chromosomal genes in Escherichia coli K-12 using PCR products. Proc Natl Acad Sci U S A 97:6640–6645.

4.	Andris-Widhopf J, Rader C, Steinberger P, Fuller R, Barbas CF. 2000. Methods for the generation of chicken monoclonal antibody fragments by phage display. J Immunol Methods 242:159–181.
