## Supplementary material for "Improvement of affinity and potency of a monoclonal antibody against *Shigella flexneri* 3a O-antigen via phage display and whole-cell in-solution panning": Table S3

| Name | Sequence (5’-3’) | Purpose |
| --- | --- | --- |
| pKD4F | GTGTAGGCTGGAGCTGCTTC | Amplify Kan cassette |
| pKD4R | CATATGAATATCCTCCTTAG | Amplify Kan cassette |
| waaL_downstreamSceIF | TGGCGGCCGCGGGAATTCGATTTAGGGATAACAGGGTAAT TTGGCCGCTGCTCAC | Amplify downstream for waaL mutagenesis |
| waaL_downstreamR | GAAGCAGCTCCAGCCTACACGTAAAAAATAAAAAAGGCTGC | Amplify downstream for waaL mutagenesis |
| waaL_upstreamF | CTAAGGAGGATATTCATATGGTCATCTTATCTCCGATG | Amplify upstream for waaL mutagenesis |
| waaL_upstreamSceIR | GCCGCGAATTCACTAGTGATATAGGGATAACAGGGTAAT GCGATTATGCTATCGCA | Amplify upstream for waaL mutagenesis |
| tolR_upstreamSceIF | TAGGGATAACAGGGTAATTTGCTTCTGCTTTAACTCGG | Amplify upstream for tolR mutagenesis |
| tolR_upstreamR | GAAGCAGCTCCAGCCTACACTAAACATCTGCGTTTCCCTTGCTTG | Amplify upstream for tolR mutagenesis |
| tolR_downstreamF | CTAAGGAGGATATTCATATGCATGGCTTACCCCTTGTTGC | Amplify downstream for tolR mutagenesis |
| tolR_downstreamSceIR | TAGGGATAACAGGGTAATTCTGGAATCGAACTCTCTCG | Amplify downstream for tolR mutagenesis |
| phage_library_Amp_F | CTCAGTGGCCCAGGCGGCC | Amplify phage display library for pCOMB3xss |
| phage_library_Amp_R | CTCAGTGGCCGGCCTGGCC | Amplify phage display library for pCOMB3XSS |
| S30G_F | GAGTATTGGTGACTACTTACA | Clone hFlex3a2_v2 variants |
| S30G_R | TGTAAGTAGTCACCAATACTC | Clone hFlex3a2_v2 variants |
| L33N_F | GACTACAATCACTGGTATCA | Clone hFlex3a2_v2 variants |
| L33N_R | TGATACCAGTGATTGTAGTC | Clone hFlex3a2_v2 variants |
| H34A_F | GACTACTTAGCGTGGTATCA | Clone hFlex3a2_v2 variants |
| H34A_R | TGATACCACGCTAAGTAGTC | Clone hFlex3a2_v2 variants |
| A51R_F | CAAATATCGTTCCCAATCCA | Clone hFlex3a2_v2 variants |
| A51R_R | TGGATTGGGAACGATATTTG | Clone hFlex3a2_v2 variants |
| N91H_F | ACTGTCAACATGGTCACAGC | Clone hFlex3a2_v2 variants |
| N91H_R | GCTGTGACCATGTTGACAGT | Clone hFlex3a2_v2 variants |
