## Supplementary figures and images for "Improvement of affinity and potency of a monoclonal antibody against *Shigella flexneri* 3a O-antigen via phage display and whole-cell in-solution panning"

### Figure S1

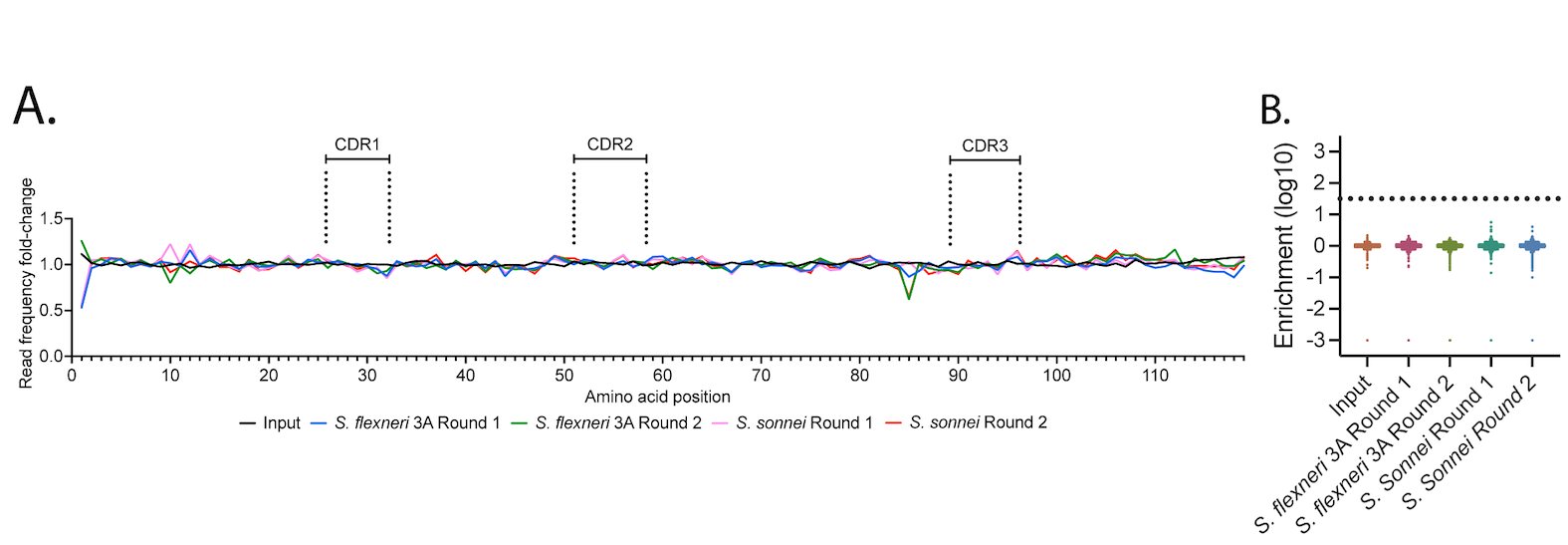

### Figure S2

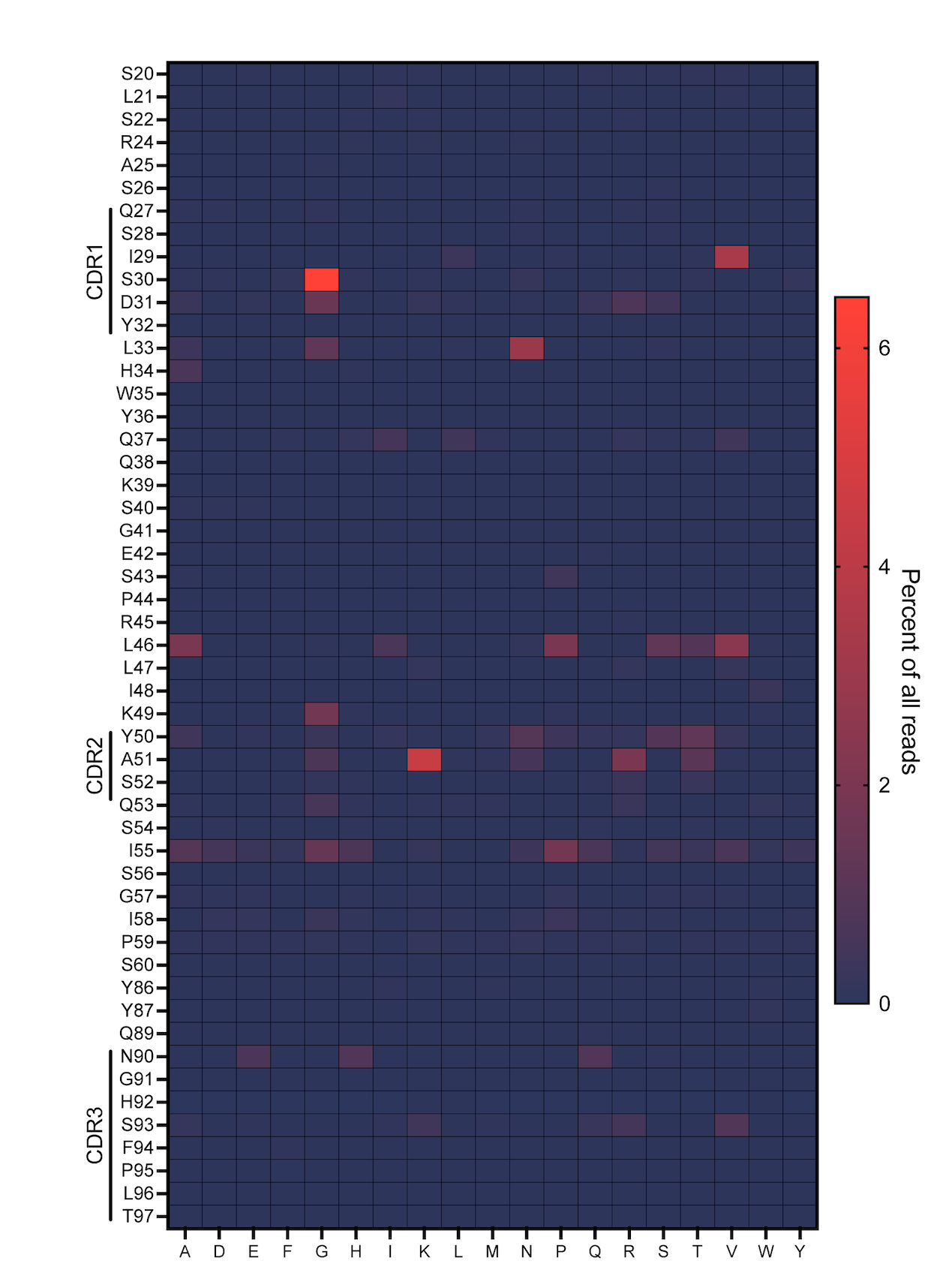

### Figure S3

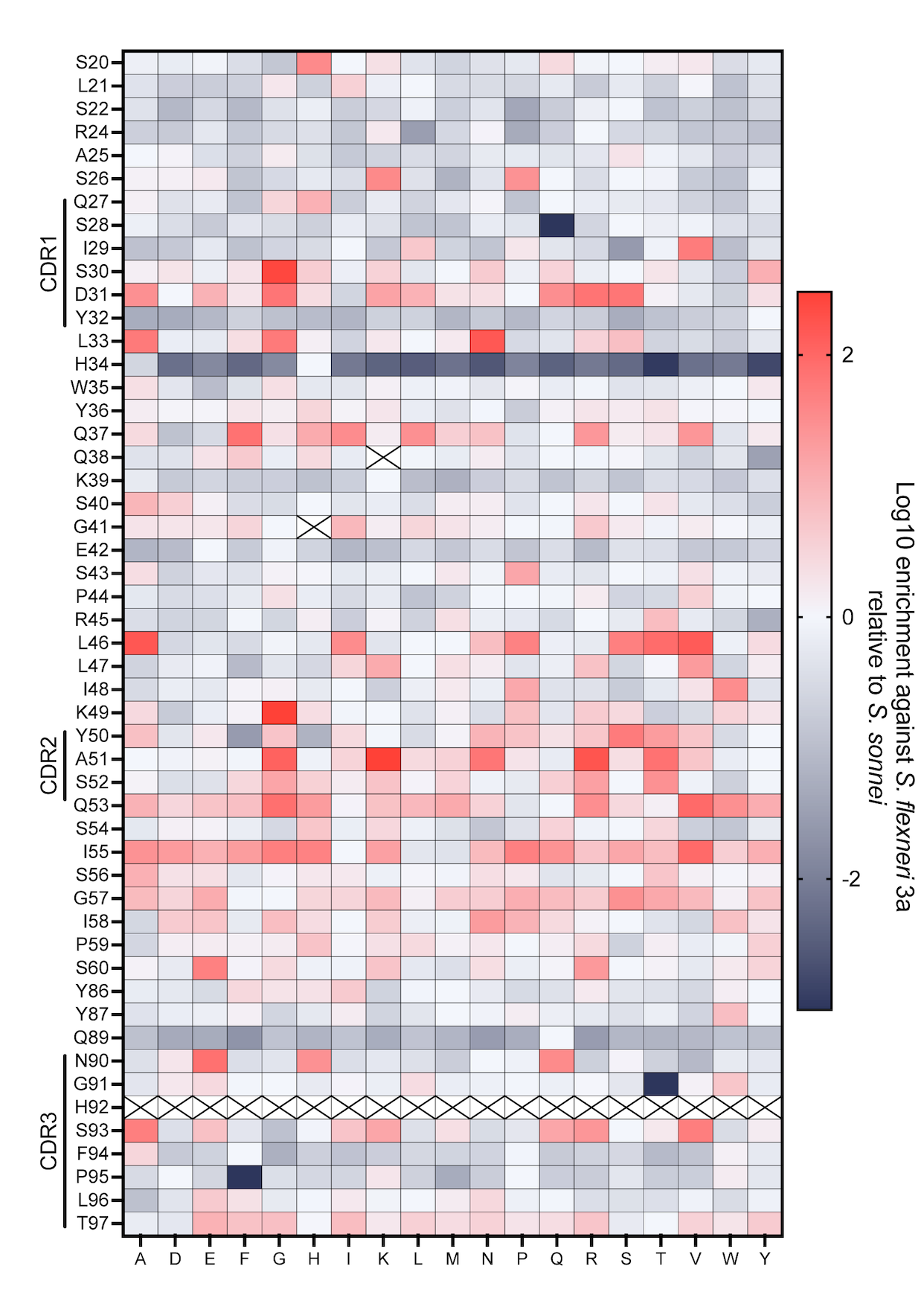

### Figure S4

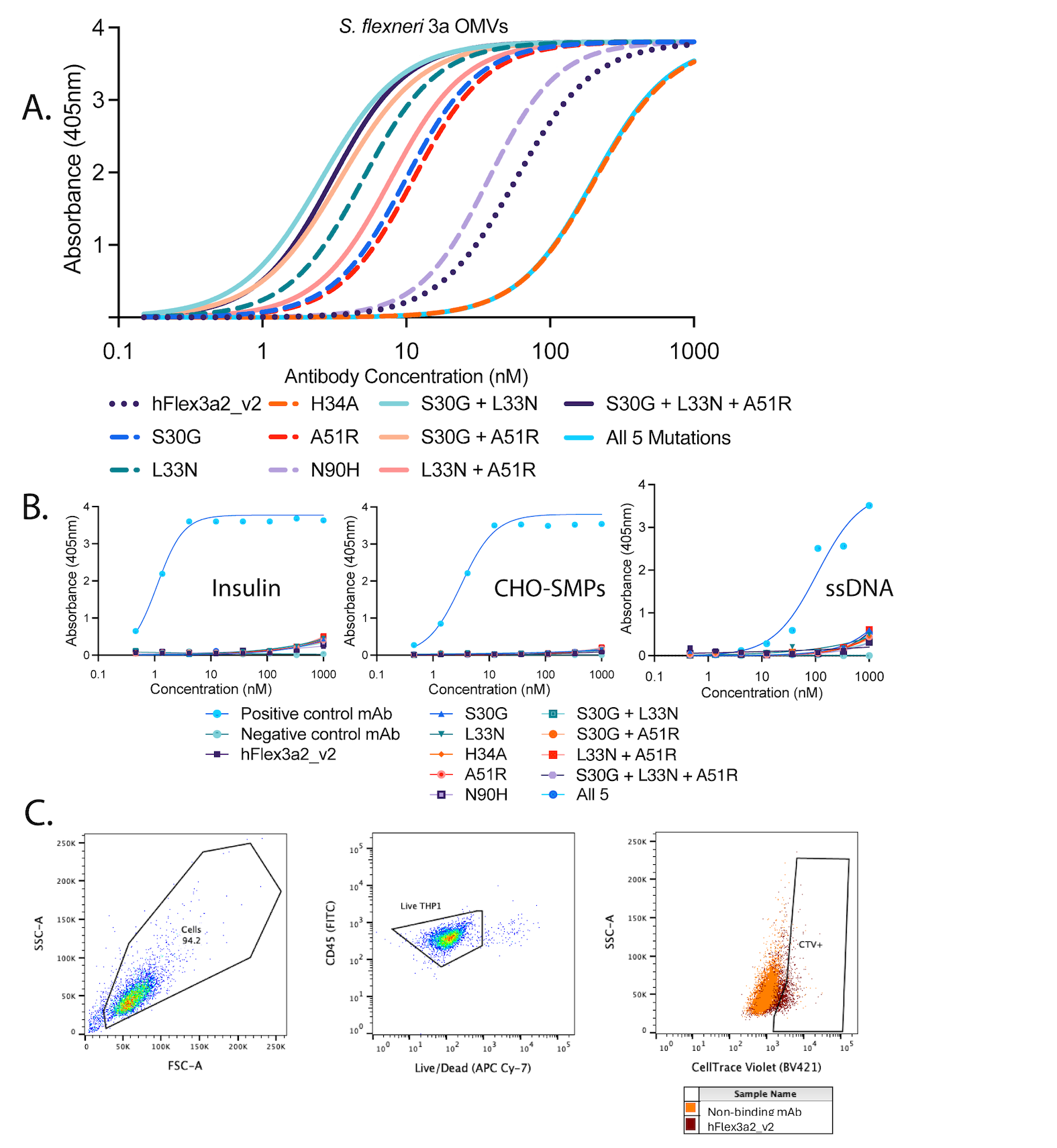

### Figure S5

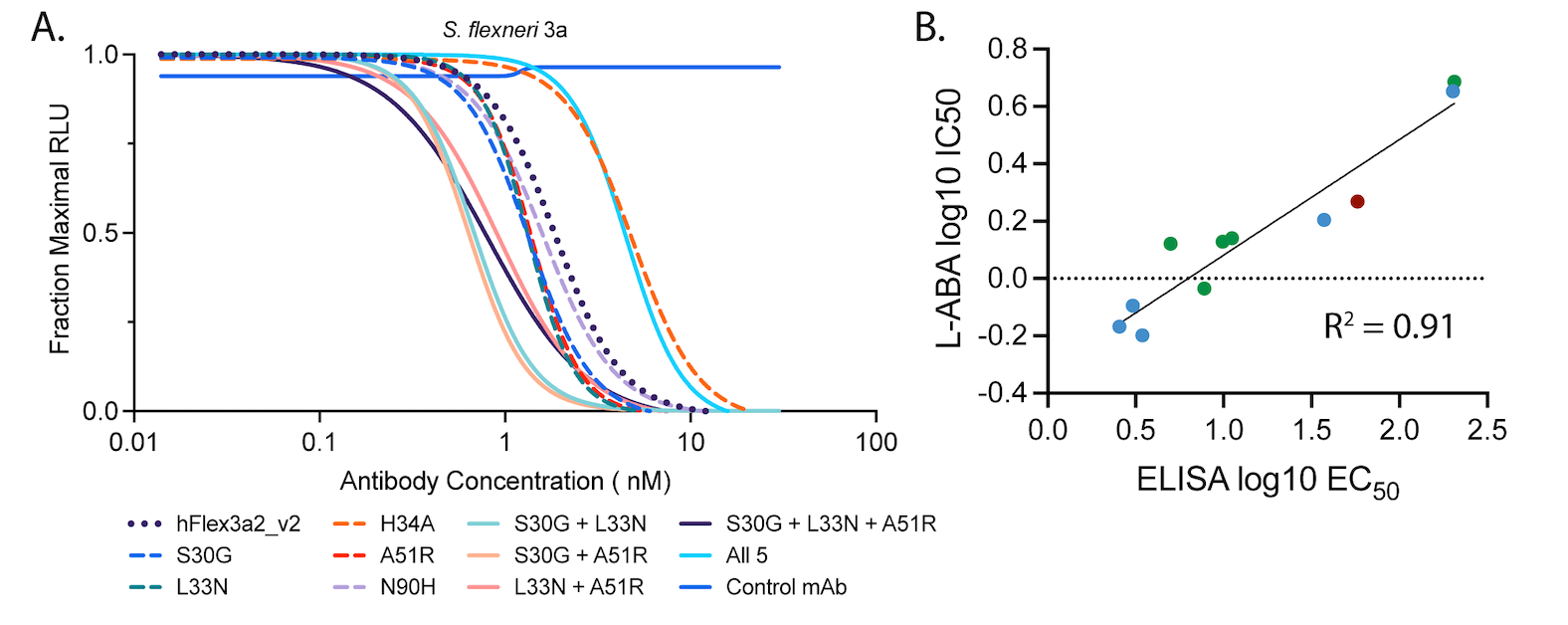

### Figure S6

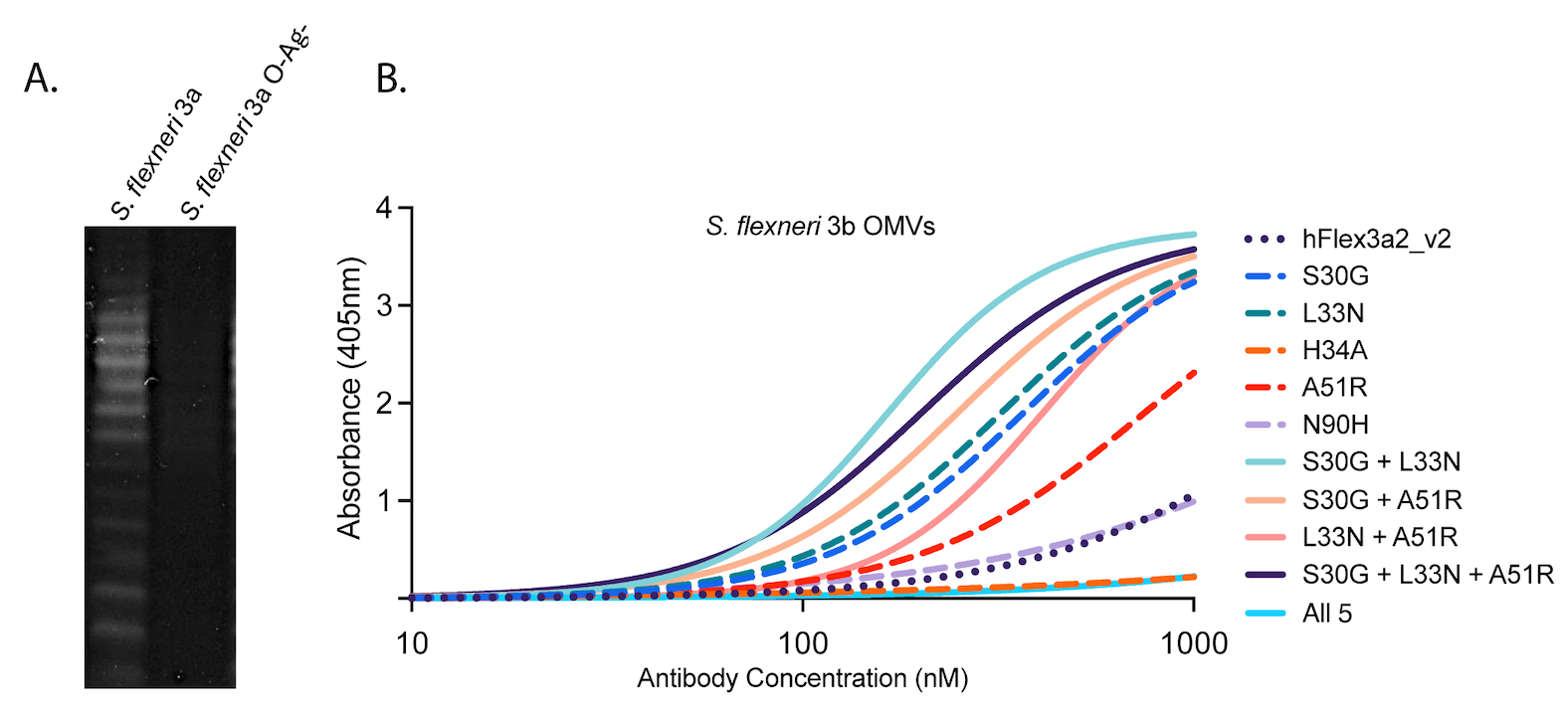
